# Evaluating the efficacy of protein quantification methods on membrane proteins

**DOI:** 10.1101/2024.04.02.587709

**Authors:** Jana Löptien, Sidney Vesting, Susanne Dobler, Shabnam Mohammadi

## Abstract

Protein quantification is an important tool for a wide range of biological applications. The most common broadscale methods include the Lowry, bicinchoninic acid (BCA), and Coomassie Bradford assays. Despite their wide applicability, the mechanisms of action imply that these methods may not be ideal for large transmembrane proteins due to the proteins’ integration in the plasma membrane. Here, we investigate this problem by assessing the efficacy and applicability of these three common protein quantification methods on a candidate transmembrane protein – the Na,K-ATPase (NKA). We compared these methods to an ELISA, which we newly developed and describe here for the quantification of NKA. The use of a relative standard curve allows this ELISA to be easily adapted to other proteins and across the animal kingdom. Our results revealed that the three conventional methods significantly underestimate the concentration of NKA compared to the ELISA. Further, by applying the protein concentrations determined by the different methods to in vitro assays, we found that variation in the resulting data was consistently low when the assay reactions were prepared based on concentrations determined from the ELISA. Thus, when target protein concentrations vary across samples, the conventional quantification methods cannot produce reliable results in downstream applications. In contrast, the ELISA we describe here consistently provides robust results.

## 1. Introduction

Many biological applications rely on protein quantification. These include biochemical assays that assess the function of proteins [1,2], the assessment of protein abundances between tissues and species [3], and the assessment of knockdown experiments [4]. The most common all-encompassing methods of protein quantification include the Lowry, bicinchoninic acid (BCA), and Coomassie Bradford assays [5–7]. In general, these methods are sensitive, simple, inexpensive, and easily reproducible [8]. Moreover, they are available as standardized commercial kits. In all three methods, total protein concentration is detected by a color signal whose intensity correlates with the quantity of total protein present in the sample and which can be measured spectrophotometrically.

Both the Lowry and BCA assays are copper-based quantification methods. They rely on the reduction of cupric ions (Cu^2+^) to cuprous ions (Cu^+^) – a process known as the biuret reaction. This reaction is facilitated by the peptide bonds of proteins and the subsequent biuret signal formed intensifies in proportion to the amount of Cu^+^ in the sample [5,6,9]. Therefore, the more peptide bonds are accessible to Cu^2+^ in the reaction, the stronger the biuret signal, and thus the higher the determined protein concentration will be.

In the Lowry assay, a strong alkaline cupric sulfate and tartrate reagent is added to the protein sample and during a short incubation, light blue tetradentate Cu^+^ complexes form (the biuret reaction). These complexes are formed between Cu^2+^ and the nitrogen atoms present in the peptide bonds of proteins. In a secondary reaction, Cu^2+^ is reduced to Cu^+^ by the protein peptide bonds (oxidation of peptide bounds) [5,9]. Following incubation, the yellow Folin-Ciocalteu reagent (phosphotungstic acid and phosphomolybdic acid mixture in phenol) is added and color strengthening occurs. The precise mechanism for color strengthening using the Folin-Ciocalteu reagent is not completely understood, but it is believed to occur when the tetradentate Cu^+^ complexes transfer electrons to the phosphomolybdic-phosphotungstic acid complexes (Cu^+^ is oxidized back to Cu^2+^ by Mo^6+^) [10]. The reduced phosphomolybdic-phosphotungstic acid complexes produced by this reaction are intensely blue in color (heteropolymolybdenum Blue (Mo^5+^)) and can be measured spectrophotometrically at around 750 nm [5]. Some specific amino acids (tryptophan, tyrosine, and cysteine) also contribute to the color strengthening by the Folin-Ciocalteu reagent [11]. Thus, proteins with higher frequency of these amino acids will produce stronger color signals.

The BCA is a modification of the Lowry assay. BCA replaces the Folin-Ciocalteu reagent to generate a method that is more tolerant to interference by commonly encountered substances present in protein samples [6]. The advantage of the BCA is that it is relatively stable under alkaline conditions, unlike the Lowry assay. Thus, BCA can be incorporated into the alkaline copper solution in a simplified one-step procedure [6,12]. At room temperature, the readily oxidizable tryptophan, tyrosine, and cysteine or cystine are the main amino acids responsible for Cu^2+^ reduction in this assay. At higher temperatures (37-60°C), Cu^2+^ is also reduced by protein peptide bonds [6,13]. However, experiments with di-, tri– and tetrapeptide proteins suggest that the extent of color formation is caused by more than the mere sum of individual color-producing functional groups in the protein and that the peptide backbone likely contributes as well [13]. In the final step of the reaction, each reduced Cu^+^ chelates with two molecules of BCA, forming a purple complex that can be measured spectrophotometrically at around 562 nm [6,14].

In contrast to the Lowry and BCA assays, the Coomassie Bradford protein assay (Bradford assay) is based on the binding of the Coomassie Brilliant Blue G-250 dye to protein, which results in protein-dye complexes [7]. The Bradford assay is the fastest and simplest of all protein quantification methods. The strong non-covalent binding of the dye to proteins is due to interactions with hydrophobic amino acids tryptophan and phenylalanine and the polar amino acid tyrosine. The binding of the neutral species to proteins also occurs due to electrostatic attraction between the negatively charged sulfonic group of dye and the positively charged guanidino group of arginine and, to a lesser degree, of the other basic amino acids, lysine and histidine. Protein-bound dye molecules shift their absorbance maximum near 615 nm [15], and the increase in absorbance around that wavelength is proportional to the number of bound dye molecules [7].

All three conventional protein quantification methods have advantages and drawbacks. One of the clear drawbacks is that they detect all proteins in a given sample and can thus only provide an estimate for specific protein concentrations when samples contain unpurified or partially purified proteins. Moreover, all three methods exhibit some degree of detection variation towards different protein types (protein-to-protein variation). This can be attributed to the different reagents’ affinities towards certain amino acids, as described above. Indeed, the determined protein concentrations can be significantly over– or underestimated depending on the amino acid composition of the proteins [16], with the Bradford assay showing approximately twice as much detection variability compared to the copper-based protein quantification methods [14]. The BCA and Lowry assays appear to have very similar detection variability between different proteins [6].

Another limitation of these conventional methods is that the proteins must be solubilized to interact with the assay reagents. This makes the methods susceptible to interference from other substances present in protein samples. For example, the Bradford assay is sensitive to surfactants and detergents (e.g., SDS or Triton X-100) at concentrations routinely used to lyse cells and solubilize membrane proteins [7]. This is because detergents also bind to proteins and compete with the dye for binding sites, thus preventing the color reaction [17]. The Lowry and BCA assays are compatible with most surfactants, even if present in the protein sample at concentrations up to 1 % (Lowry assay) or 5 % (BCA assay) [14]. On the other hand, the Bradford assay is less impacted by substances that reduce copper (e.g., dithiothreitol or ß-mercaptoethanol) or copper-chelating agents (e.g., EDTA), which are normally used to stabilize proteins in solution [7,12]. Buffer solutions that change the pH of the reagents cause problems for all three conventional protein quantification methods. In general, the Bradford assay is more tolerant towards interfering substances, like buffer salts, than the Lowry and BCA assays [18], and the BCA assay is less prone to disruption than the Lowry assay [6,14]. In addition, the Bradford reagent is highly acidic, thus, proteins with poor acid-solubility cannot be assayed with this reagent.

Another challenge faced by these conventional methods, which we focus on here, is when proteins are embedded in lipid membranes. Transmembrane proteins are largely embedded in the cell’s hydrophobic lipid bilayer, with one or more membrane-spanning regions that have hydrophobic amino acid side chains distributed on their surface. Often, these transmembrane proteins have additional hydrophilic domains located on the extra– and intracellular regions. Membrane proteins have proven to be difficult to purify owing to their integration in the plasma membrane, partially hydrophobic surfaces, flexibility, and lack of stability [19]. Moreover, some transmembrane proteins are only present in low abundance, and it can be difficult to obtain enough protein for accurate quantification [20]. The Bradford assay has already been shown to underestimate the protein concentration of membrane-containing fractions due to the significant portion of membrane proteins being embedded in the cell membrane and thus not accessible to the dye molecules [21]. Therefore, experimental work with such proteins has often relied on relative protein quantification by a combination of Bradford assays and western blots [22–26]. While these methods can be sufficient for estimating protein concentrations across samples, they lack the precision needed for producing robust measurements, especially when target protein concentrations vary across biological replicates. Moreover, western blots are labor-intensive, and not ideal for explicit quantification.

Enzyme-linked immunosorbent assays (ELISAs) present a solution for quantifying large transmembrane proteins and overcome some of the limitations and drawbacks associated with the conventional methods. ELISAs are used to quantify specific target proteins in a sample. Thus, they overcome the specificity limitations that conventional protein quantification methods face with heterogenous protein samples. This specificity is achieved in ELISAs through antibody-antigen interactions and an enzyme-mediated color reaction to detect the presence of the target protein (antigen). This interaction allows the quantification of small amounts of a specific target protein in a heterogenous sample [27,28]. However, due to their specificity, commercial ELISAs for less commonly studied proteins and for proteins from non-model organism are hard to come by and cost prohibitive.

ELISAs follow the same basic principles [28,29]. The first step is adhering or immobilizing the target protein from the liquid phase directly onto the surface of 96-well plate wells. Following this, unsaturated surface-binding sites are blocked by non-reacting protein, such as bovine serum albumin (BSA). Next, the target protein is detected either directly (enzyme-labeled primary antibody that binds specifically to the target protein) or indirectly (such as enzyme-labeled secondary antibody that detects the primary antibody-target protein interaction). In the final step, substrates for the enzyme are introduced and enzymatically converted to a chromogenic, fluorescent, or luminescent signal. The intensity of the signal directly correlates with the amount of target protein present in the sample.

Three types of ELISAs have been developed: sandwich, direct, and indirect. A sandwich ELISA is the most precise method because the target protein is bound between two primary antibodies (capture antibody and detection antibody), each detecting a different epitope of the target protein. However, this method requires a lot of tailoring for each protein type and is thus not easily applicable across a broad range of proteins. A direct ELISA uses a single primary antibody, which also serves as the detection antibody, allowing the direct detection of the target protein. An indirect ELISA relies on a secondary antibody to detect the primary antibody. A standard secondary antibody can be applied to different primary antibodies, and the primary antibody does not require any labeling. The indirect ELISA is thus the most easily applicable method. Although a variety of commercial ELISA kits are available for different proteins, they are expensive, specific for only a narrow range of protein sources (e.g., rats, mice, humans) and can be ineffective depending on the protein isolation methods.

To address the challenge of accurate transmembrane protein quantification, we compare the three conventional methods (Lowry, BCA, and Bradford) and evaluate their performance in detecting and quantifying transmembrane proteins in comparison to an indirect ELISA, which we newly developed and validate here. We use Na, K-ATPase (NKA) as our model protein. This transmembrane protein has been studied extensively in our working group and we have established a high-throughput expression system to produce NKA from a wide range of taxa. The smallest functional unit of the NKA consists of a dimer of a large α-subunit with 10 transmembrane helices and a glycosylated β-subunit of one transmembrane helix [30,31].

The ELISA developed for NKA in this paper makes use of a commercially available primary antibody, which binds universally across the animal kingdom. Attempts to produce a standard from commercially available purified NKA proved ineffective. Therefore, we additionally developed a method for producing relative standards by lyophilizing an aliquot of the protein being measured. These specifications allow our ELISA to be adapted to any protein type and source.

## 2. Material and Methods

All key resources and reagents used in this study are listed in detail in Table S1.

### Acquiring expression vectors

The majority of our protein samples were produced by expression in cell culture. This allowed us to sample from a broader range of taxa. We used six pFastBac Dual plasmids, which have two cloning sites for the simultaneous expression of two proteins, containing the NKA α1– and β1-subunit genes from six different animal species. The species included the brown rat (*Rattus norvegicus*), common ostrich (*Struthio camelus*), gold tegu (*Tupinambis teguixin*), large milkweed bug (*Oncopeltus fasciatus*), Miranda’s white-lipped frog (*Leptodactylus macrosternum*), and red-necked keelback snake (*Rhabdophis subminiatus*). We selected these species because we wanted to sample as broad a range of animals as possible. All vertebrate vectors were originally produced by Mohammadi et al., [1,26,32] and the large milkweed bug vector was produced by S. Dalla [32]. Each vector had the α1-subunit gene (ATP1A1 for vertebrates, ATPα1C for the large milkweed bug) under the control of the P_PH_ promoter and the β-subunit gene (ATP1B1 for vertebrates, β1 for the large milkweed bug) under the p_10_ promoter. Accession numbers for the sequences and the Addgene plasmid numbers associated with these plasmid constructs can be found in Table S2.

### Bacmid production

The six recombinant plasmids were transposed into competent DH10Bac *Escherichia coli* cells and grown on plates containing gentamycin, tetracycline, kanamycin, Isopropyl β-D-1– thiogalactopyranoside (IPTG), X-Gal. DH10Bac cells contain a bacmid and a helper plasmid. The transposed donor plasmid segment containing the inserted ATP1A1 and ATP1B1 genes, along with a gentamicin resistance gene, recombines at the *lac*Z alpha peptide gene on the bacmid, disrupting its reading frame. The resulting colonies are white when grown on plates containing IPTG and X-Gal, whereas those with intact *lacZ* proteins, i.e., failed recombination, are blue. Following transposition, two white colonies were picked from each plate (per NKA construct) and incubated in liquid culture containing gentamycin, tetracycline, and kanamycin to amplify the recombinant bacmids.

The amplified bacmid DNA was isolated from *E. coli* cells via alkaline lysis. Briefly, the *E. coli* cells were lysed with a strong alkaline solution consisting of 200 mM NaOH and 1% SDS. Cell debris, SDS, and genomic DNA was removed by centrifugation, and the resulting bacmid DNA was pelleted and washed with 70% ethanol. The pellet was resuspended in purified water and the resulting bacmid DNA was analyzed by PCR and agarose gel electrophoresis to verify that the transposition of the genes of interest to the bacmids was successful (Fig. S1). The primers used for this analysis were pUC/M13 FWD 5’– CCCAGTCACGACGTTGTAAAACG-3’ and pUC/M13 RVS 5’-AGCGGATAACAATTTCACACAGG-3’.

### Generation of recombinant baculoviruses and expression of NKA proteins in Sf9 cells

Sf9 cells (immortalized ovary cells from the moth *Spodoptera frugiperda*) were used as the host for recombinant baculovirus production and protein expression. Sf9 cells are advantageous for NKA expression because they express very little of the protein endogenously [33,34].

We followed the modified one-step protocol described by Scholz and Suppmann [35]. For baculovirus production, 8×10^6^ Sf9 cells in 10 mL Insect-Xpress medium with 30 µg/mL gentamycin were transfected with 10 µg purified bacmid DNA using PEI MAX. We repeated this procedure three times to produce three biological replicates (A, B, C) per NKA construct. After 5 days of incubation at 27°C and 110 rpm, the Sf9 cells were pelleted by centrifugation at 500 x g for 5 min and the P0 viruses in the supernatant were harvested. Freshly seeded Sf9 cells (50×10^6^ cells in 50 mL media) were subsequently infected with the P0 viruses (500 µL virus per biological replicate). Following three days of incubation at 27°C and 110 rpm, the Sf9 cells were pelleted by centrifugation at 1650 x g for 10 min and stored at –80°C until membrane isolation. Additionally, uninfected Sf9 cells (i.e., without NKA expression) were also cultured (25×10^6^ cells in 50 mL media) and pelleted for use as a negative control.

### Membrane isolation from Sf9 cells

The frozen Sf9 cell pellets were resuspended in 15 mL of homogenization buffer (0.25 M sucrose, 2 mM EDTA, 25 mM HEPES; adjusted to pH 7.0 with Tris HCl) on ice. The cell suspensions were then sonicated at 60 W (Sonopuls 2070) for three 45 s intervals on ice to lyse the cells and disrupt the cell membranes. Next, the cell suspensions were centrifuged for 30 min at 10,000 x g and 4°C (Centrifuge 5840 R) to pellet cell debris. The supernatant was collected and further centrifuged for 60 min at 146,000 x g and 4°C (Ultra-Centrifuge L-80) to pellet the cell membranes. The supernatant was removed, and the pelleted membranes were washed twice and resuspended in cooled HPLC grade water (1 mL per biological replicate), then stored at –20°C.

### Preparation of tissue samples

Brain and kidney tissue was dissected from three freshly frozen black rats purchased commercially from frostmaus.de. Nervous tissue (brains, prothoracic and central ganglia) was also dissected out of 14 freshly frozen large milkweed bugs and pooled into one sample. The bugs were acquired from a colony reared at University of Hamburg [36]. All tissue samples were homogenized on ice with a glass grinder in 500 µL HPLC grade water. Nervous tissue samples were lyophilized overnight (12-24 h) at 0.1 mbar and –50°C in a freeze dryer. After lyophilization, the freeze-dried samples were reconstituted in 500 µL (large milkweed bug) or 1 mL (black rat) cooled HPLC grade water. The reconstituted samples were then sonicated in an ice bath for 10 min (Omni Sonic Ruptor 400, Pulser 90, Power 90%) and centrifuged for 10 min at 5000 x g and 4°C (Centrifuge 5840 R) to precipitate debris. The resulting supernatant was stored at –20°C. Membranes from kidney samples were isolated as described above for Sf9 cells. Nervous tissue is known to have very high levels of NKA expression compared to other tissues [37–40]. Therefore, we deemed it unnecessary to concentrate the proteins by isolating membranes for this tissue.

### Positive control

A lyophilized sample of commercially available, purified NKA from porcine cerebral cortex (0.3 units/mg protein; Table S1) was reconstituted to a stock concentration of 10 mg/mL in HPLC grade water.

### Testing the effect of lyophilization on antibody detection

We wanted to prepare relative standards for our ELISA by using the dry weight of lyophilized protein samples containing NKA from different species. We therefore first tested whether lyophilizing NKA would cause a reduction in their detection by antibodies. For this comparison we used a 300 µL aliquot of membrane isolated samples derived from Sf9 cell expression (Table 1), which consisted of 100 µL of each replicate (A-C) pooled together. For common ostrich and Miranda’s white-lipped frog, 162.5 µL of replicates A and B were pooled because the C replicates were contaminated during the preparation process. The samples were dried overnight (12-24 h) at 0.1 mbar and –50°C in a freeze dryer. Afterwards, the samples were centrifuged shortly and their weights calculated by subtracting the weight of the empty tubes. The freeze-dried samples were then reconstituted in 200-300 µL cooled HPLC grade water. An aliquot was taken from each sample for the downstream production of relative standards.

**Table 1.** List of protein samples used in this study. Protein samples derived from tissue are highlighted in grey, all others were expressed in Sf9 cells.

| Order | Species | NKA Source | Used for |
| --- | --- | --- | --- |
| Mammalia | *Rattus norvegicus*<br>(brown rat) | Expressed in Sf9 cells | (1) testing effects of lyophilization<br>(2) relative NKA standard production<br>(3) comparing protein quantification methods<br>(4) comparing applicability of quantification methods |
| Mammalia | *Rattus rattus*<br>(black rat) | Brain tissue preparation | (3) comparing protein quantification methods<br>(4) comparing applicability of quantification methods |
| Mammalia | *Rattus rattus*<br>(black rat) | Kidney tissue preparation | (3) comparing protein quantification methods<br>(4) comparing applicability of quantification methods |
| Mammalia | *Sus domesticus*<br>(porcine) | Commercially available,<br>lyophilized cerebral cortex<br>NKA isolate | (2) NKA standard production<br>(3) comparing protein quantification methods<br>– **positive control** |
| Aves | *Struthio camelus*<br>(common ostrich) | Expressed in Sf9 cells | (1) testing effects of lyophilization<br>(2) relative NKA standard production<br>(3) comparing protein quantification methods<br>(4) comparing applicability of quantification methods |
| Squamata | *Tupinambis teguixin*<br>(gold tegu) | Expressed in Sf9 cells | (1) testing effects of lyophilization<br>(2) relative NKA standard production<br>(3) comparing protein quantification methods<br>(4) comparing applicability of quantification methods |
| Squamata | *Rhabdophis subminiatus*<br>(red-necked keelback<br>snake) | Expressed in Sf9 cells | (1) testing effects of lyophilization<br>(2) relative NKA standard production<br>(3) comparing protein quantification methods<br>(4) comparing applicability of quantification methods |
| Amphibia | *Leptodactylus macrosternum*<br>(Miranda's white-lipped<br>frog) | Expressed in Sf9 cells | (1) testing effects of lyophilization<br>(2) relative NKA standard production<br>(3) comparing protein quantification methods |
| Insecta | *Oncopeltus fasciatus*<br>(large milkweed bug) | Expressed in Sf9 cells | (1) testing effects of lyophilization<br>(2) relative NKA standard production<br>(3) comparing protein quantification methods<br>(4) comparing applicability of quantification methods |
| Insecta | *Oncopeltus fasciatus*<br>(large milkweed bug) | Brain and ganglia tissue<br>preparation | (3) comparing protein quantification methods |
| Insecta | untransfected Sf9 cells<br>(ovary cells from<br>*Spodoptera frugiperda*) | Sf9 cells (contain very little<br>to no endogenous NKAs) | (1) testing effects of lyophilization<br>(3) comparing protein quantification methods<br>– **negative control** |

For the western blot analysis, 2.5 μL of non-lyophilized and reconstituted lyophilized samples were solubilized in 8 μL 4x Laemmli buffer (62.5 mM Tris HCl pH 6.8; 2 % SDS; 10 % glycerol; 5 % 2-mercaptoethanol; 0.001 % Bromophenol Blue) and 21.5 μL Millipore water. The solubilized samples were then separated on an SDS gel containing 10% acrylamide. Subsequently, the separated proteins were blotted from the gel onto nitrocellulose membrane. To block unsaturated binding sites on the membranes after blotting, they were incubated at room temperature in 1x BlueBlock PF on a plate tipper at medium tipping speed for 1 h. After blocking, the membranes were transferred into a 50 mL Falcon tube and incubated with 2 mL of the primary antibody (Table S1) at a concentration of 0.88 μg/mL in 1x BlueBlock PF overnight at 4°C on a tube rotator. Next, the membranes were washed three times with 1x BlueBlock PF. The membranes were then incubated with 2 mL of the secondary antibody (Table S1) at a concentration of 8 μg/mL in 1x BlueBlock PF for 1 h at room temperature on the tube rotator. Next, the membranes were washed three times with 1x BlueBlock PF and two additional times with 0.05 M Tris HCl for 5 min each. The antibody complexes were stained by moving the membranes into a glass petri dish and adding the detection enzyme substrate (0.035 % H_2_O_2_, and 0.01 % 4-Chloro-1-naphthol in 0.05 M, pH 7.5 Tris HCl). Following a 10 min incubation, the staining reaction was stopped by washing the membrane with deionized water. The resultant membranes were scanned (Canon 9000F Mark II). These samples were also compared on our ELISA (described below) and the results from both the western blot and ELISA were statistically compared (described below). See Table S3 for raw data.

### Protein assays

All the protein samples we produced were quantified by Lowry, BCA, Bradford, and ELISA. The commercial porcine sample was used as a positive control, and the protein sample derived from untransfected Sf9 cells was used as a negative control. All four assays were run on aliquots that were not previously freeze-thawed.

### Lowry protein assay

We used Thermo Fisher Scientific’s Modified Lowry protein assay kit (Table S1). First, a standard curve was prepared with bovine serum albumin (BSA) (1500, 1000, 750, 500, 250, 125, 25, 5, 1, and 0 μg/mL)

in Millipore water. The protein samples were diluted 1:10 in Millipore water, except for porcine and large milkweed bug tissue samples, which were not diluted because low protein content was expected in these samples. 40 μL of the standards and protein samples were pipetted in duplicate (two technical replicates) into 96-well polystyrene flat-well plate wells. 200 µL of Lowry reagent was added to each well, mixed on a microplate shaker for 30 seconds, and then incubated at room temperature for 10 min. Following incubation, 20 μL of freshly prepared 1x Folin-Ciocalteu reagent was added to each well and mixed on the microplate shaker for 30 seconds. Following a 30 min room temperature incubation, the absorbance of standards and protein samples was measured at 655 nm using a microplate absorbance reader (Bio-Rad Model 680). See Table S4 for raw Lowry assay data.

### Bicinchoninic acid (BCA) protein assay

We used Thermo Fisher Scientific’s Bicinchoninic acid protein assay kit (Table S1). First, a standard curve was prepared with BSA (1500, 1000, 750, 500, 250, 125, 25, 5, 1, and 0 μg/mL) in Millipore water. The protein samples were diluted 1:10 in Millipore water, except for porcine and large milkweed bug tissue samples, which were not diluted. 25 μL of the standards and protein samples were pipetted in duplicate (two technical replicates) into 96-well polystyrene flat-well plate wells. 200 µL of freshly prepared, clear, green BCA working reagent was added to each well, and mixed on a microplate shaker for 30 seconds. Following a 30 min incubation at 37°C and subsequent cooling of the plate to room temperature, the absorbance of standards and protein samples was measured at 550 nm using a microplate absorbance reader (Bio-Rad Model 680). See Table S5 for raw BCA assay data.

### Coomassie Bradford protein assay

For the Bradford assay, we prepared a standard curve with BSA (10, 8, 6, 4, 2, and 0 μg/mL) in Millipore water. The protein samples derived from Sf9 cells and rat brain tissue were diluted 1:2000 in Millipore water, whereas porcine and large milkweed bug tissue samples were diluted 1:200 (see methods above). Both the standards and protein samples were then diluted 1:2 with Coomassie Bradford reagent and pipetted into polystyrene cuvettes in duplicate (two technical replicates). After incubating at room temperature for 10 mins, the absorbance of standards and protein samples was measured at 595 nm using a cuvette spectrophotometer (Ultrospec 2100 pro). See Table S6 for raw Bradford assay data.

### Relative NKA standard preparation for the ELISA

For each animal species to be assayed, a relative NKA standard was prepared from the lyophilized membrane isolates derived from Sf9 cells (Fig. 1). The concentration of each standard was calculated based on their lyophilized weights (described above). These concentrations were used to create relative NKA standard stock aliquots of 1 mg/mL in ELISA coating buffer (0.1 M NaHCO_3_ in PBS, adjusted to pH 9.5 with KOH; Table S7) for each animal species.

**Figure 1.**
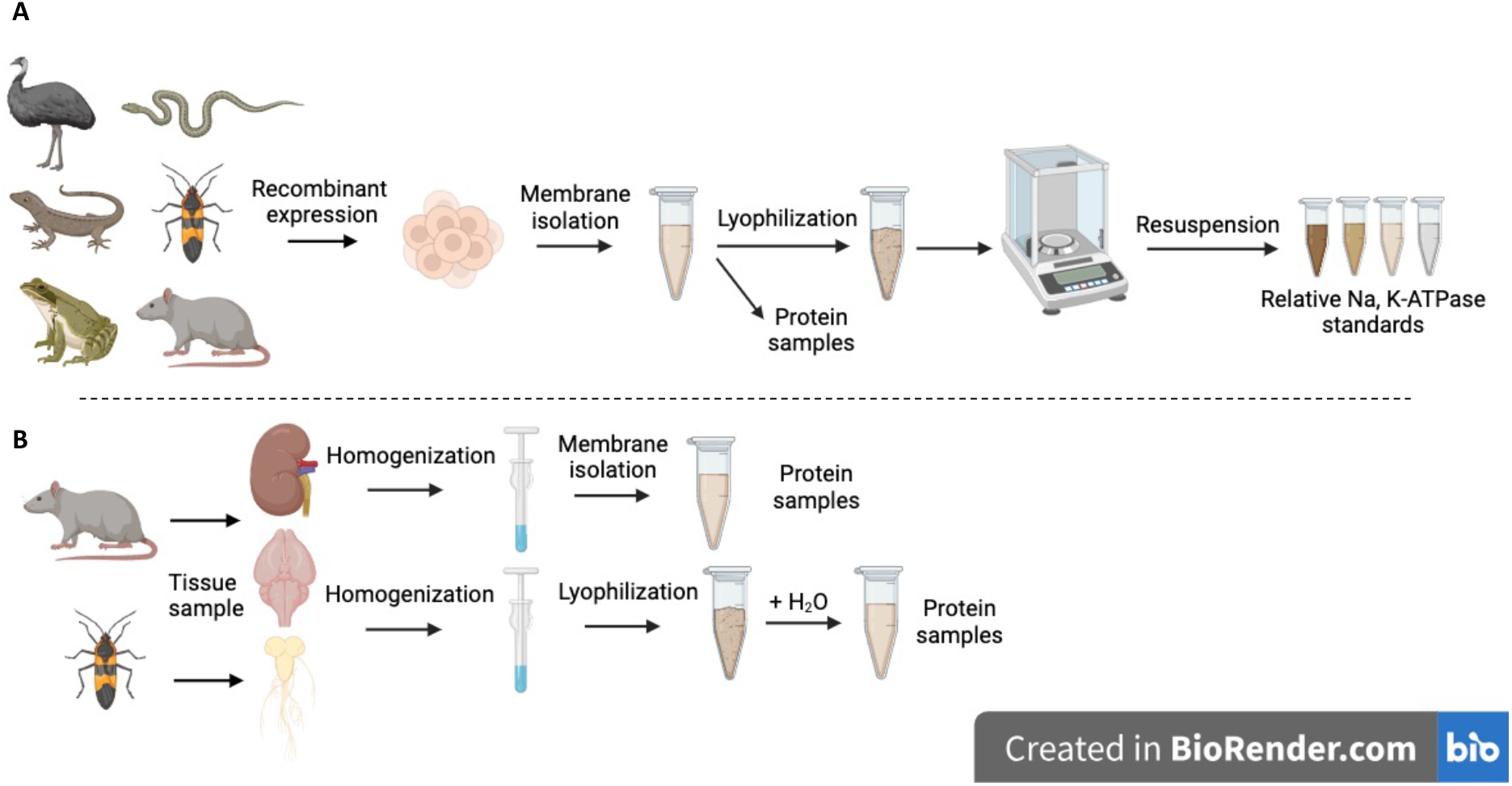
Schematic diagram illustrating the isolation of proteins from (A) recombinant expression in Sf9 cells and (B) from tissues. Nervous tissue samples were isolated by homogenization, followed by centrifugation to pellet waste and lyophilization, whereas all other samples were membrane isolated. Relative Na, K-ATPase (NKA) standards were subsequently prepared from a subset of membrane isolates from recombinantly expressed Sf9 cells. All resulting protein samples were used in all four quantification assays, while the relative NKA standards were produced specifically for the ELISA. Figure created with BioRender.com.

### Indirect ELISA to quantify NKA proteins

Prior to running an ELISA, the protein samples were diluted 1:2000 in the coating buffer (Table S7). Replicates A-C of common ostrich and large milkweed bug were diluted 1:6000 and 1:4000, respectively, because higher NKA concentrations were expected in these samples based on the western blots. The 1 mg/mL relative NKA standard stocks were sonicated in an ice bath for two 2.5-min intervals (Omni Sonic Ruptor 400, Pulser 90, Power 90%) and afterwards the standards were diluted in coating buffer to the following concentrations: 60, 30, 20, 15, 10, 7.5, 5, 3.75, 2.5, 1.875, 1.25, 0.9375, 0.625, and 0 μg/mL. All ELISAs were run on aliquots that were not previously freeze-thawed.

First, 100 μL of protein samples and relative NKA standards were added to 96-well polystyrene flat-well plates in duplicate (two technical replicates). Additionally, 100 μL of a negative control using only coating buffer without any protein was added to the plates. This negative control is equal to the 0 μg/mL standard. After adding samples and standards to the wells, the plates were sealed with Parafilm® and incubated overnight at 4°C (12-18 h).

The next day, unbound leftovers were discarded and plates dried by tapping them several times face-down on a paper towel. Each well was washed by adding 400 μL of washing buffer (0.05 % Tween® 20 in PBS; Table S7), letting it sit for 10 s, then discarding fluids as described above. This washing procedure was repeated to a total of five times. Next, unsaturated surface-binding sites were blocked by adding 200 μL of blocking buffer (1 % BSA, 0.02 % Tween® 20 in PBS; Table S7) to each well, sealing the plates with Parafilm®, and incubating at room temperature for 2 h. Following blocking, the wells were washed as described above.

We used the universal monoclonal primary antibody α5 to detect the NKA α-subunit (α5-antibody; Table S1). The antigen species for this antibody is chicken and the host species is mouse. The α5-antibody binds a cytosolic epitope on the α-subunit of all NKA isoforms across the animal kingdom. 50 μL of primary antibody solution containing 2 μg/mL α5-antibody in blocking buffer was added to each well, and left to incubate, sealed with Parafilm®, for 1 h at room temperature (overnight (12-18 h) at 4°C is also possible).

To remove unbound primary antibodies, the wells were washed as described above. Next, we used a goat-anti-mouse polyclonal secondary antibody conjugated with horseradish peroxidase (HRP-antibody; Table S1) to detect the primary antibody. The host species for the secondary antibody is goat. 50 μL of secondary antibody solution containing 5 μg/mL HRP-antibody in blocking buffer was added to each well, and left to incubate, sealed with Parafilm®, for 1 h at room temperature.

Following the secondary antibody incubation, each well was washed seven times as described above to maximally reduce background noise caused by unbound HRP-antibody. The colorimetric reaction of HRP was triggered by adding 100 μL 3,3’,5,5’-tetramethylbenzidine (TMB) to each well, and allowing color development at room temperature for 10 min. The colorimetric reaction was stopped by adding 100 μL stopping solution (0.5 M H2SO4) to each well. Finally, the absorbance of standards and protein samples was measured at 490 nm using a microplate absorbance reader (Bio-Rad iMark^TM^). See Fig. 2 for a schematic diagram of the ELISA methods and Table S7 for buffer recipes.

**Figure 2.**
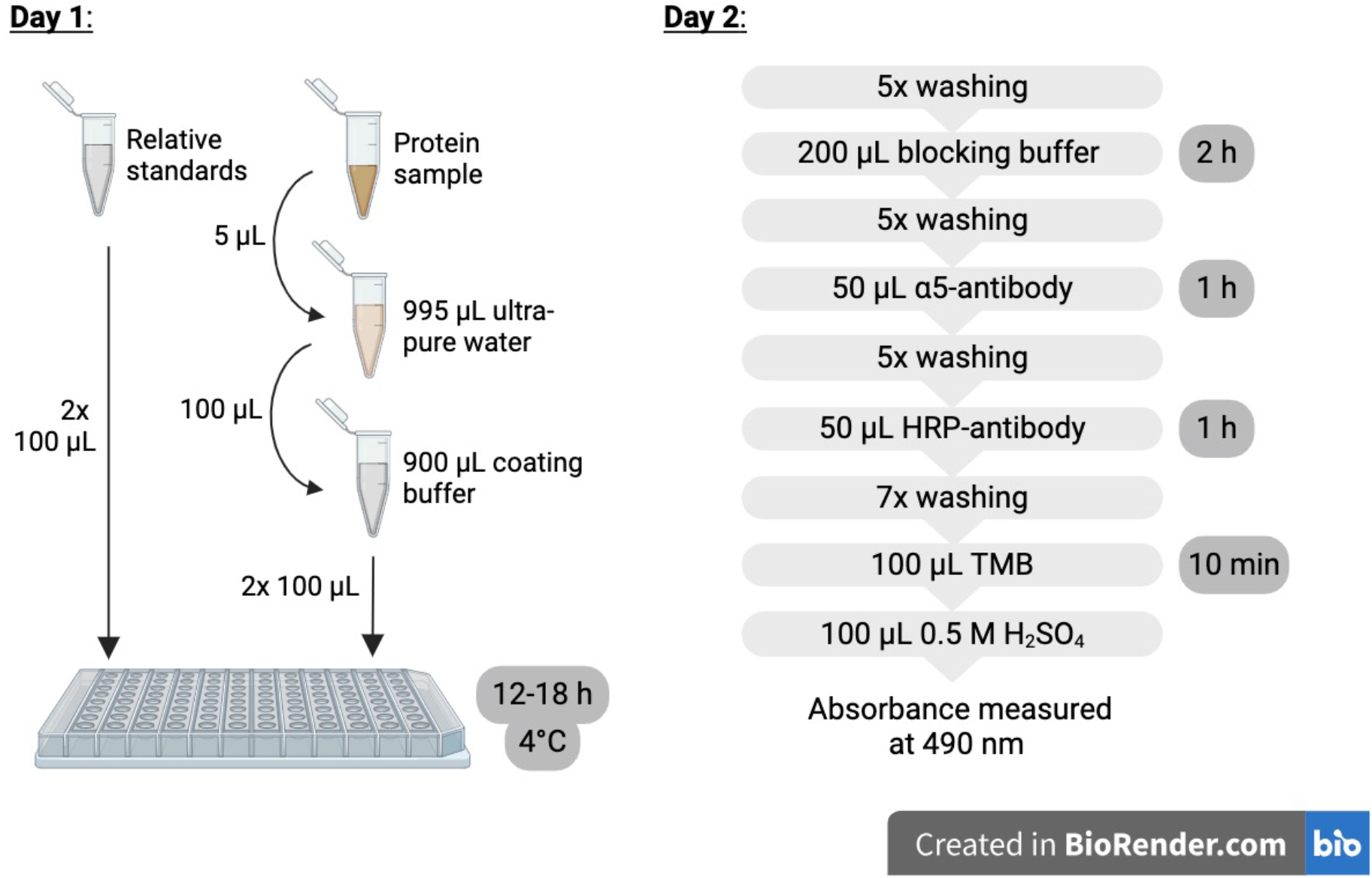
Schematic diagram illustrating the two-day ELISA procedure. Figure created with BioRender.com.

### Negative controls

Several negative controls were included to validate the functionality of this ELISA. First, the primary antibody (α5-antibody) was omitted to confirm that there is no non-specific binding of the secondary HRP-antibody. Next, the HRP-antibody was omitted to confirm that TMB and the stopping solution do not stain non-specifically (i.e., without HRP). For these negative controls, a random sample from our Sf9 expressed proteins was used (gold tegu replicate A). Then, both protein and antibodies were omitted to confirm that there is no background noise caused by the buffers. Finally, membrane isolate of uninfected Sf9 cells was run in the ELISA to confirm that the antibodies are not binding non-specifically to other membrane proteins. See Table S8 for raw negative control data.

### Calculation of protein concentrations

For all protein quantification methods, the average absorbance of the two blank standard replicates (0 µg/mL) was subtracted from each absorbance measurement. Then, the average values of the two technical replicates were calculated. The average absorbance values of each standard were used for a linear trendline-fitting. For the ELISA, either the entire range up to 60 μg/mL (red-necked keelback snake, commercial porcine NKA) or concentrations ranging from 0 to 10, 20, or 30 µg/mL were used, depending on the extent of the linear range of the standard curves. The resulting linear equations were used to calculate the protein concentrations of the samples. See Table S9 for raw ELISA data.

### Testing the robustness of protein quantification methods with in vitro NKA activity assays

To examine the robustness of the determined protein or NKA concentrations, the variation across biological replicates of in vitro NKA activity assays were compared. Only animal species with three or more biological replicates were tested in the NKA activity assays. Thus, Miranda’s white-lipped frog and large milkweed bug tissue were excluded from the assays. Replicate C of Miranda’s white-lipped frog encountered infection problems during transfection whereas the number of large milkweed bugs we dissected was only sufficient to produce only one biological replicate for the tissue. For each assay, 100 μg of protein was used and this was calculated based on concentrations determined by the four different protein quantification methods. The 100 μg of protein was pipetted in duplicate (two technical replicates) into two wells on a 96-well polystyrene flat-well plate. The first well contained stabilizing buffer to measure total ATPase activity and the second well contained inhibiting buffer (lacking KCl and including 10^-2^ M ouabain) to block NKAs and measure background ATPase activity. See assay buffer formulas in Petschenka et al. [41]. The proteins were incubated at 37°C and 200 rpms for 10 min on a microplate shaker.

Next, ATP (10 mM Tris-ATP, pH 6.5) was added to each well and the proteins were incubated again at 37°C and 200 rpms for 20 min. The activity of NKAs was determined by quantification of inorganic phosphate (P_i_) released from enzymatically hydrolyzed ATP. Reaction P_i_ levels were measured according to the procedure described by Petschenka et al. [41]. The reactions were stopped with 10 % SDS and then stained with a solution consisting of Taussky-Shorr color reagent [42], which turned the reactions blue in proportion to their P_i_ concentration. After 10 min of staining, the absorbance of each well was measured at 655 nm using a microplate absorbance reader (Bio-Rad Model 680). Average absorbance values of the two technical replicates were used for subsequent calculations.

The averages of two technical replicates were translated to mM Pi based on a standard curve of KH_2_PO_4_ (0 – 1.2 mM P_i_) that was run on the same plate. Background ATPase Pi concentrations were subtracted to obtain NKA ATPase P_i_ concentrations [41]. The P_i_ concentrations released from 100 μg of protein were then converted to nmol P_i_/(mg protein*min).

### Statistical analyses

To test whether lyophilization alters NKA detection by antibodies, we compared the distribution of ELISA absorbance values between lyophilized and non-lyophilized samples with a Wilcoxon signed-rank test for paired samples. In addition, we compared the densities of the western blot’s NKA α-subunit bands (∼110 kDa) between the two treatments using ImageJ 1.53k to draw plots from the signal strength and size of the western blot bands. From the resulting plots, the areas under the curves were calculated [43]. We then compared the distribution of the areas under the curves between lyophilized and non-lyophilized samples with a Wilcoxon signed-rank test for paired samples. For both analyses, the values for all samples were pooled together based on the treatment.

We used a Friedman rank sum test for paired samples to compare the NKA detection capabilities of the four different quantification methods. We first compared the concentration values between Sf9 and tissue derived samples and found a significant difference (Wilcoxon paired test; p < 0.0001). We thus continued by analyzing Sf9 and tissue derived samples separately. The positive control (commercial porcine sample), the negative control (uninfected Sf9 cells), and replicate C of Miranda’s white-lipped frog (which was infected by bacteria in cell culture) were excluded from this analysis. For the remaining, determined protein concentrations of different samples were pooled for each quantification method respectively. A two-sided Dunn’s post hoc test (R package “rstatix” [44]) was used to identify the quantification methods that significantly differ from one another in the distribution of their determined protein concentrations. Since multiple pairwise comparisons were performed simultaneously in this test, the Bonferroni adjustment method was used for controlling the family-wise error rate.

To evaluate the robustness of the determined NKA concentrations across the four methods, a one-way analysis of variance (ANOVA) was used to compare the mean variation (scatter) in NKA activities (specified by standard deviation of ATPase activity) across the three biological replicates (A, B, C). The NKA activity assays were based on 100 μg of protein calculated from concentrations determined by the four quantification methods. In the ANOVA, animal species was considered a random effect. Also, standard deviations and coefficients of variation of different animal species were pooled for each quantification method respectively. A power analysis (R package “pwr” [45]) revealed high power (>99 %) of the ANOVA to detect true effects or differences in the data. The black rat brain and kidney tissue samples were excluded from the analysis, because the amount of P_i_ released from these samples was beyond the linear range of the P_i_ standard curve. Despite running several additional rounds with more diluted samples, we were never able to capture an activity range within the P_i_ standards before using up the tissue samples.

All statistical analyses were performed using R 4.1.1 software (R Core Team, 2021) and a significance level of 0.05 was applied. See Supplementary Dataset 1 for data table used for statistical comparisons.

## 3. Results

### Lyophilization does not affect NKA detection by antibodies

To test whether lyophilization affects the detection of NKA by the ELISA antibodies, we compared Sf9 membrane isolates that were lyophilized to their non-lyophilized counterparts on a western blot (Fig. 3A) and in the ELISA. We analyzed the distribution of the western blot band densities and found no significant difference between lyophilized and non-lyophilized samples (Wilcoxon signed-rank paired test; p = 0.81; W = 16; n = 7; Fig. 3B). The ELISA corroborated the same result (Wilcoxon signed-rank paired test; p = 0.08; W = 25; n = 7; Fig. 3B); showing no significant difference in the distribution of the absorbance values between the two treatments. These results confirm that lyophilization of Sf9 cell membrane isolates does not significantly diminish the detectability of the NKA proteins by the antibodies used in the ELISA.

**Figure 3.**
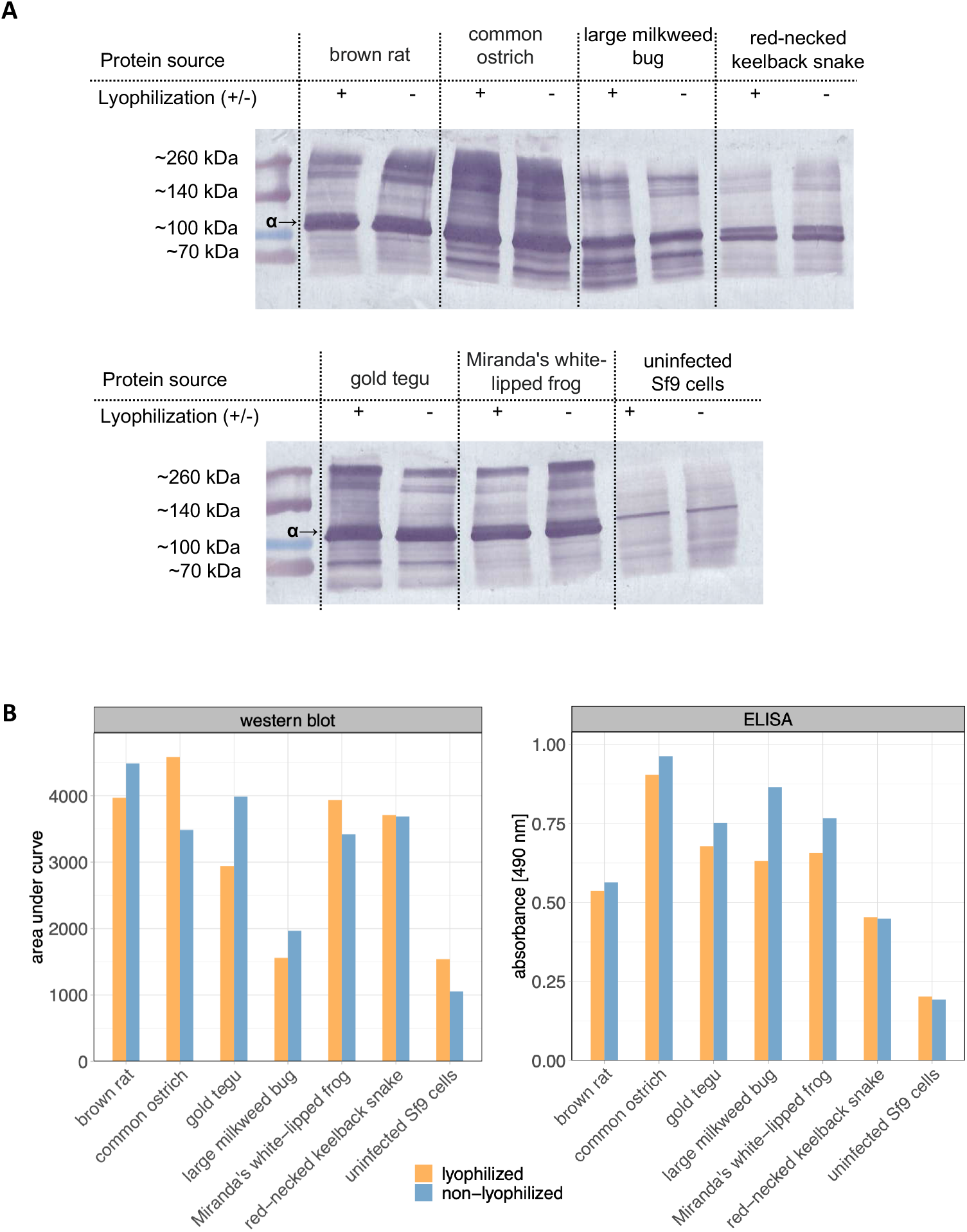
(A) Western blot membrane scans show lyophilized and non-lyophilized Sf9 cell membrane isolates run on two gels. The 110 kDa line corresponds to the NKA’s α-subunit, which is marked by an arrow. Because the α5 primary antibody is specific only to the NKA α-subunit, additional bands and smearing are most likely caused by higher order complexes and high protein load. The + and – indicate which bands are lyophilized (+) and which are not (–). (B) Comparison of detection intensity between lyophilized and non-lyophilized Sf9 membrane isolates in the western blot and ELISA analyses. Densities of the western blot’s NKA α-subunit bands were determined by the areas under the curves in ImageJ (Fig. S3; Table S3).

### Relative NKA standard curves vary between different animal species

Because the α5 antibody can have different binding affinities to NKA from different animals, we produced a standard curve for each species. These standards were prepared from NKA expressed in Sf9 cells for consistency and reproducibility. In addition, we prepared standards from commercial porcine NKA. We found that most standard curves reached saturation well before the higher concentration range (0 μg/mL to 60 μg/mL; Fig. 4A). Omitting the saturated range allowed us to capture the linear range needed for appropriate concentration calculations (Fig. 4B). The red-necked keelback snake standard was an exception to this pattern, for which a linear curve fit over the entire range of standards (Fig. 4A) – this is likely due to it having very low levels of NKA expression and thus lower amounts of NKA per mg lyophilized membrane isolate (Fig. S2). The standards prepared from commercial porcine NKA exhibited similarly low detectability by the α5 antibody. This was, however, surprising because the commercial porcine sample contains only NKA in membrane fragments and thus each mg is pure NKA. Because of the high variation, having standards tailored to each animal group ensures that the concentrations of samples are not over– or underestimated in the ELISA.

**Figure 4.**
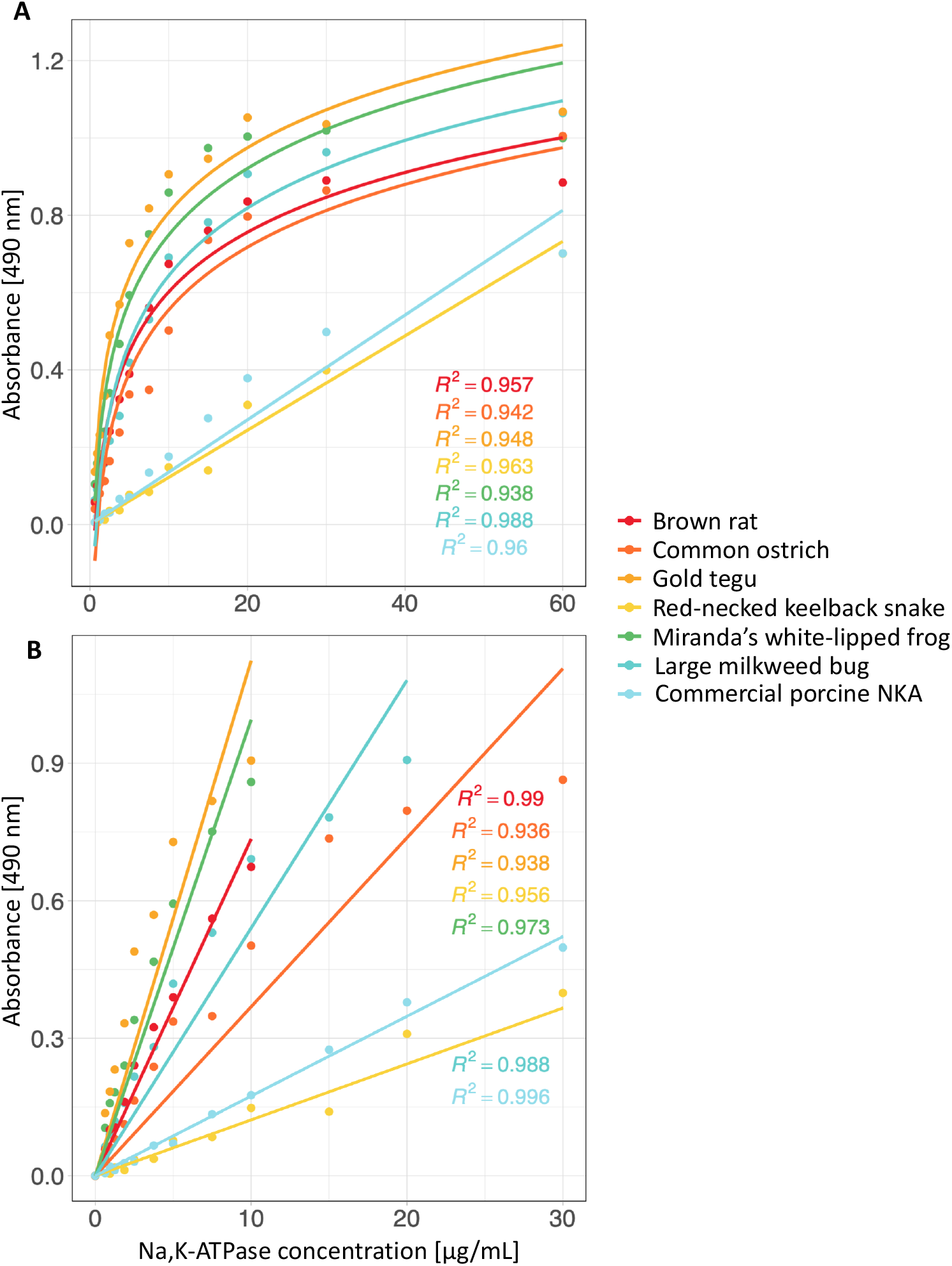
Comparison of standard curves generated from Sf9 cell membrane isolates containing recombinantly expressed Na, K-ATPase (NKA) from different animal species. Additionally, a standard curve prepared from purified commercial porcine NKA is included. (A) The full range of the relative NKA standards (0 to 60 μg/mL) reached saturation mid-way for most animal species. (B) Omitting the saturated range allowed us to capture the linear range of the standard curves, which were subsequently used for calculating NKA concentrations.

### NKA detection is minimal to absent in negative controls

To assess the specificity of the antibodies used in the ELISA, several negative controls, which excluded different components of the ELISA, were tested. We set the detection threshold to 0.034, which is the average absorbance value of the reagents used in the ELISA (containing neither protein nor antibody). We found that the absorbance of the negative control containing no protein (i.e., membrane isolate) remained within this detection threshold (Fig. 5; Table S8). The negative control containing no HRP-conjugated secondary antibody fell slightly above this threshold. The negative control containing no primary α5-antibody minimally exceeded the detection threshold, which indicates that there is slight background noise caused by the HRP-conjugated secondary antibody. This is likely caused by unbound secondary antibody that remains in the well after repeated washing. In addition to these negative controls, we tested the NKA-specificity of the primary α5-antibody using membrane isolates of untransfected Sf9 cells, which express very little NKA endogenously [33,34]. There was some detection of NKAs in these membranes. This is expected because these cells express a small level of these proteins endogenously. In addition, untransfected cells have higher cell proliferation and thus membranes isolated from these samples will have a higher density of membranes than those isolated from transfected cells. Nevertheless, the detection of NKAs in samples derived from transfected Sf9 cells exceeded untransfected cells by more than two-fold (Fig. 5). Together, these results confirm that the specificity of the ELISA for NKA is suitable.

**Figure 5.**
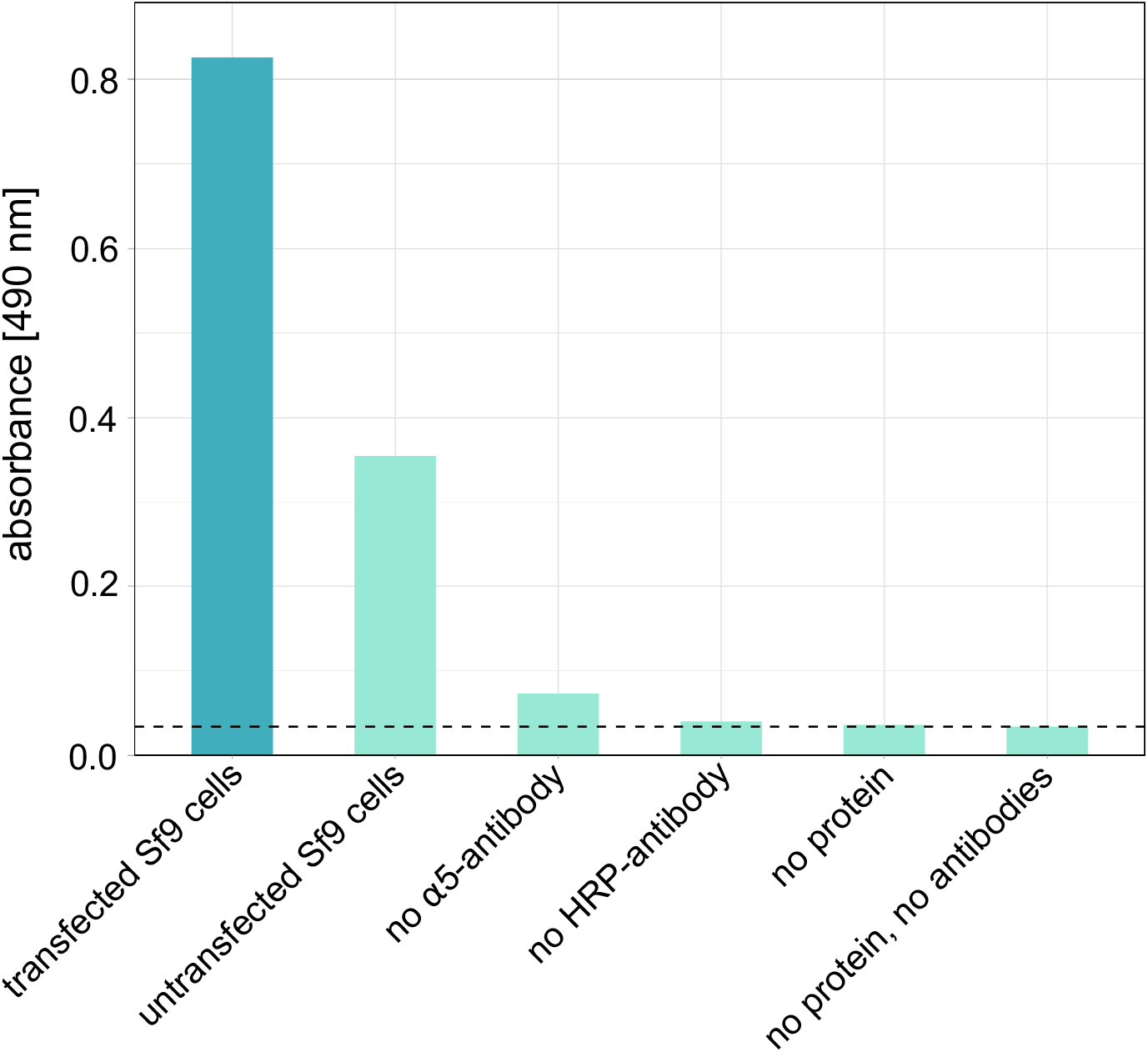
Comparison of absorbance values between a protein sample derived from transfected Sf9 cells (positive control; blue) and various negative controls (light turquoise). The positive control consisted of a gold tegu (replicate A) sample. This standard was subsequently used in the negative controls that contained protein (i.e., no α5-antibody and no HRP-antibody). The dashed line shows the detection threshold of “no protein, no antibodies”. Each bar represents the average of two technical replicates from one biological replicate.

### Conventional protein quantification methods cannot sufficiently quantify NKA

Our comparisons between protein quantification methods revealed several notable patterns (Fig. 6A). In general, there was strong interspecies variation in protein concentrations among samples quantified by ELISA, which ranged between 1.39 to 145.89 mg/mL. This variation was not as pronounced across the samples quantified by conventional methods, which all-together ranged between 0.55 to 18.27 mg/mL (Supplementary Dataset 1). The absence of stark variation among the conventional quantification methods strongly points towards their inability to robustly quantify NKA, and possibly all transmembrane proteins.

**Figure 6.**
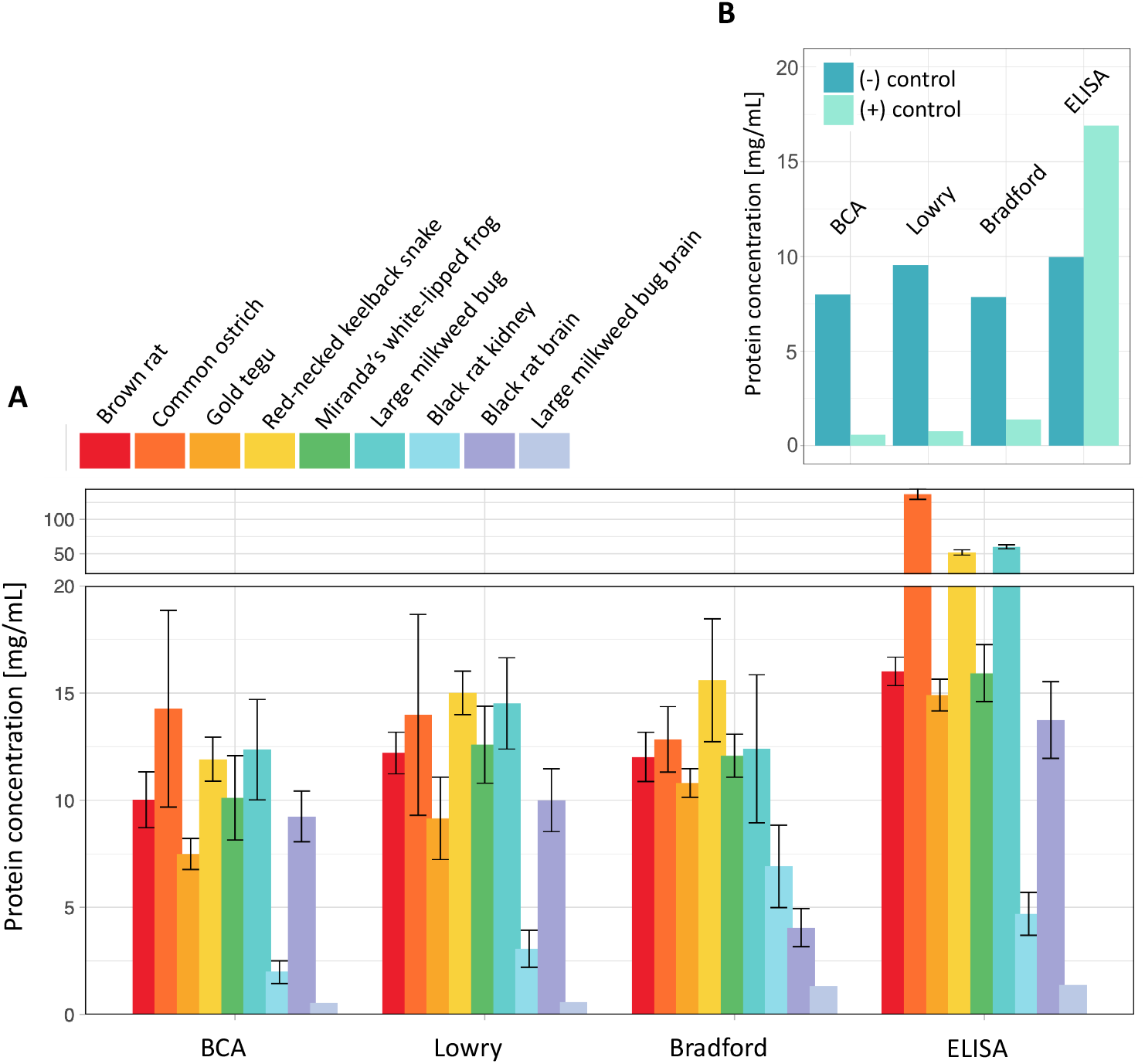
(A) Protein concentrations of recombinantly expressed and tissue-isolated NKAs. Bars represent the mean and standard error of three biological replicates, with the exception of Miranda’s white-lipped frog and large milkweed bug brain, which consist of two and one replicates, respectively). (B) Protein concentrations of the negative control (membrane isolates of untransfected Sf9 cells) and positive control (commercial porcine sample) determined by the four quantification methods reveals that the Lowry, BCA, and Bradford cannot sufficiently detect NKA.

Importantly, the positive control consisting of purified porcine NKA revealed that all three conventional quantification methods determined protein concentrations much lower (<1.5 mg/mL) than its actual concentration (10 mg/mL) (Fig. 6B; Supplementary Dataset 1). In contrast, the ELISA determined a concentration 16.91 mg/mL, which is not only higher than the conventional methods but also exceeds the predicted concentration based on lyophilized weight. All four quantification methods determined approximately the same concentrations in the negative control consisting of membrane isolates from untransfected Sf9 cells (Fig. 6B). Since untransfected cells have higher viability, the rather high detected protein concentrations reflect the higher overall protein content resulting from higher cell density. Taken together, these results suggest that the three conventional protein quantification methods significantly underestimate the concentration of NKA in a given sample, and likely other large transmembrane proteins.

To assess the NKA detection capabilities of the four different protein quantification methods, we pooled the determined concentrations of each method and statistically compared the resulting concentration distributions with the Friedman rank sum test for paired samples (Fig. 7A-B). We found that the protein concentration distributions for recombinantly expressed proteins were significantly different between the four methods (Friedman rank sum paired test; p < 0.001; χ2 = 36.53; df = 3; n = 15; Fig. 7A), with the ELISA having determined significantly higher protein concentrations compared to all three conventional methods (Dunn’s post-hoc test; Table S10). In contrast, the three conventional methods did not significantly differ in their determined protein concentrations. Remarkably, the wide range of NKA concentration differences detected by the ELISA was undetectable by the conventional methods, which is clearly illustrated by the stark difference in box blots (Fig. 7A). This indicates that using a transmembrane protein standard in place of the conventional BSA in the conventional methods would not be effective. We also found a marginally significant difference in protein concentration distributions for tissue-isolated samples (p = 0.006; χ2 = 12.6; df = 3; n = 9; Fig. 7B). However, a post-hoc Dunn’s test could not detect significant differences between any of the comparison pairs (Table S10). We did not observe as wide a range of concentration differences in the ELISA on tissue samples. This is likely due to tissue samples only covering two species (black rat and large milkweed bug).

**Figure 7.**
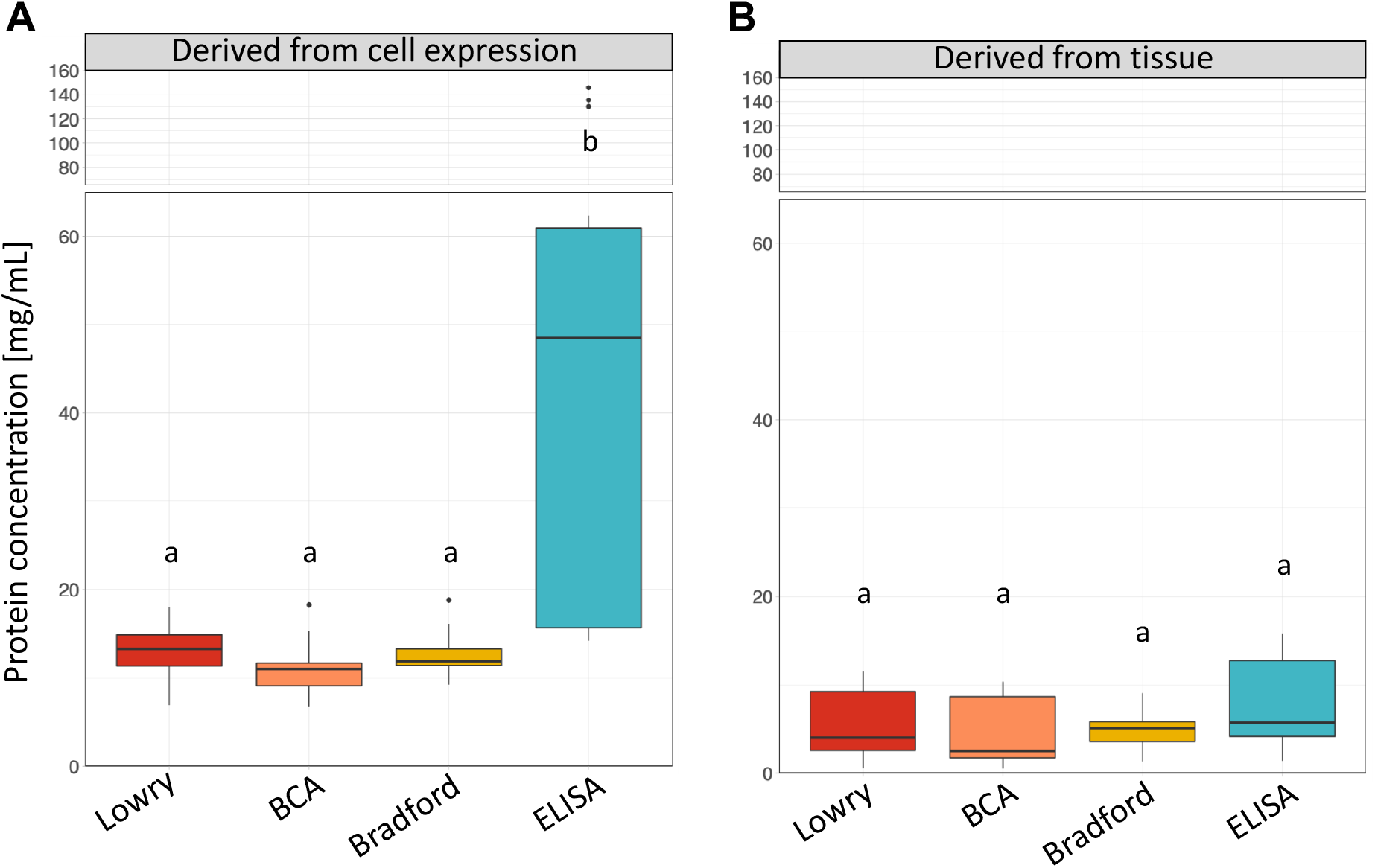
Comparison of protein concentrations determined by three conventional protein quantification methods and the ELISA from (A) recombinantly expressed NKA and (B) tissue-isolated proteins. Determined protein concentrations of different samples were pooled for each method. The boxes show the 25% and 75% quantiles of the median (thick line), the whiskers represent the maximum or minimum values without outliers. Outliers (filled-in circles) are protein concentrations greater or less than 1.5 times the interquartile range. Sample sizes for all four methods: n = 17 (samples derived from Sf9 cells) and n = 9 (nervous tissue and kidney isolates). Different letters indicate significant differences in the distribution of determined protein concentrations at p < 0.05 (Dunn’s test).

### Quantification with ELISA consistently produces more robust downstream ATPase assay data

To test whether the differences in protein quantification accuracy we uncovered affect downstream in vitro functional assays, ATPase activity assays were run based on protein concentrations determined by the four quantification methods. We tested whether the resulting variation in NKA ATPase activities (specified by standard deviation of NKA activity) significantly differs between the four quantification methods.

We did not detect a significant effect of protein quantification method on the mean standard deviation of NKA activities (ANOVA; p = 0.186; F = 1.882; n = 5; Fig. 8A). Although not statistically detectable, the variation in NKA activities of all ELISA samples is consistently low compared to those of the conventional methods, suggesting that the ELISA provides robust NKA quantification. Western blots of the recombinantly expressed NKA show that expression levels were even across biological replicates (Fig. S2). This indicates that overall protein content would be even within samples, and thus reduce variation in activity based on concentrations determined by the conventional methods.

**Figure 8.**
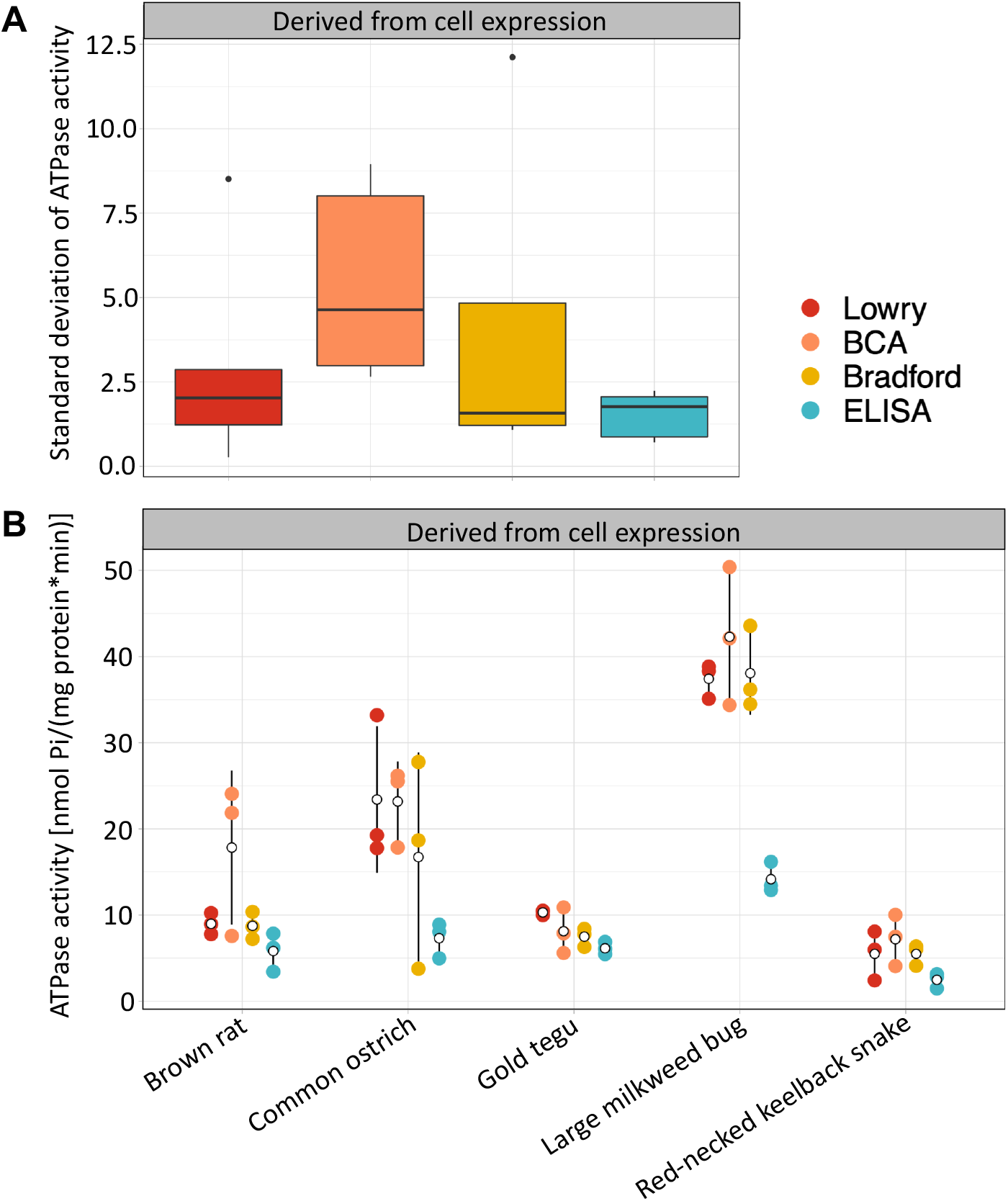
Comparison of the standard deviations (SDs) of Na, K-ATPase (NKA) ATPase activities measured based on protein concentrations determined by the four different protein quantification methods. (A) Pooled means showed no significant differences in SDs between the four methods. The box plots show the 25% and 75% quantiles of the median (thick line) and the maximum or minimum values without outliers (whiskers). Outliers (filled-in circles) are ATPase activities greater or less than 1.5 times the interquartile range. The sample size for each protein quantification method is n = 15. (B) Comparison of the variation in NKA activity of protein content calculated from concentrations determined by the four different protein quantification methods. Means and standard deviations of the three biological replicates are shown as hollow black circles and lines. Raw data points are shown as filled-in circles. Samples isolated from tissues were excluded from this analysis because activity levels fell beyond the linear range of standards.

Of note, activity measurements of the large milkweed bug NKA resulted in stark differences between the ELISA and conventional methods (Fig. 8B). The high activity associated with the conventional methods is due to them detecting very low concentrations of the milkweed bug proteins, resulting in more protein being added to the activity assay reaction, and subsequently higher measured activity. In contrast, the ELISA detected rather high NKA concentrations in the large milkweed bug, which resulted in less protein being used for the activity assays, and thus lower activity measurements. This pattern can be seen across samples but is most pronounced in the milkweed bug.

In contrast to the Sf9 derived samples, the NKA activities of tissues derived samples deviate from this pattern, showing high variation across all four quantification methods. Almost all the activity values of tissue samples fell far beyond the linear range of the P_i_ standard curve. Thus, we could not make accurate activity estimates for most of these samples. For this reason, they were excluded from the ANOVA analysis.

## 4. Discussion

The overarching aim of this paper was to find a reliable method for quantifying large transmembrane proteins. We focused on the NKA as a model and we developed an ELISA that can be easily adapted across protein types and sources (i.e., species). We assessed the specificity of our ELISA in comparisons to commonly used conventional quantification methods (Lowry, BCA, and Bradford) and found that all conventional methods are unsuitable for quantifying NKA. If the conventional methods, which should detect all proteins in a given sample, quantify transmembrane proteins efficiently, then the protein

concentrations determined by these methods should be significantly higher than those determined by the NKA-specific ELISA. Contrary to this expectation, we found that the ELISA consistently determined higher protein concentrations in all the protein samples. Notably, the concentration of the positive control, which consisted of pure NKA in membrane fragments, was grossly underestimated by the conventional quantification methods—to the point where it was almost completely undetected (Fig. 6B). These outcomes are a clear indication that the Lowry, BCA, and Bradford assays cannot detect transmembrane proteins. It is also important to point out that the ELISA also strongly underestimated the concentration of the pure porcine NKA (Fig. 6B). This commercially available protein is purified from porcine nervous tissue. According to the manufacturer, the primary α5-antibody, which we use in the ELISA, is capable of binding to a cytosolic epitope on the NKA α-subunit of all isoforms across the animal kingdom. However, it is possible that the α5-antibody binds more strongly to isoform α1 (ATP1A1), which we expressed in Sf9 cells, than to α3 (ATP1A3), which is the main isoform expressed in nervous tissue [46]. To confirm this hypothesis, it would be prudent to test the antibody’s ability to detect α1 and α3 isoforms expressed and isolated from Sf9 cells.

There are several possible explanations for why the conventional quantification methods are inefficient at detecting transmembrane proteins. We rule out interfering substances, such as detergents, as a likely explanation for the inefficiency of the conventional methods because we used HPLC grade water for the resuspension of isolated proteins. We can also rule out glycosylation as a possible source of interference. Whereas glycosylation can lead to over-(Lowry and BCA) or underestimation (Bradford) of protein concentrations [47] and the NKA β-subunit is highly glycolyzed, our results consistently show underestimation of proteins by the conventional methods.

The most likely explanation has to do with the fact that the main detection molecules of the conventional protein quantification methods, CBB and Cu^2+^, are not able to penetrate the hydrophobic cell membrane [21]. Isolated transmembrane proteins are embedded in membrane fragments and therefore the embedded portion of the protein is not accessible to CBB and Cu^2+^ molecules, which bind to specific amino acids on the protein surface. This would consequently result in underestimated protein concentrations, and even more so considering that the protein standards for the conventional methods consist of non-membrane proteins. Although penetrating the cell membrane should be less of a barrier for the Lowry method because its alkaline reaction can disrupt cell membranes through hydrolysis, the results did not show higher detectability by this method compared to the other two (Fig. 7). In contrast, the α5-antibody of the ELISA binds to an epitope on the NKA α-subunit located outside the cell membrane. Therefore, the detection pathway of the ELISA is not hindered by the membrane and the resulting concentration measurement reflects a reliable estimate of the number of NKA molecules in the sample. A possible alternative would be to produce the relative standards as described for our ELISA and use them in conjunction with one of the conventional methods. However, the very low detectability of our positive control by these methods casts doubts on whether a reliable standard curve can be obtained with transmembrane proteins.

One unexpected outcome of these experiments is that all quantification methods detected lower protein concentrations in the tissue-derived samples compared to the recombinantly expressed Sf9 cell-derived samples. It is possible that the lowered detection of proteins in tissue sample is due to differences in the membrane composition, or co– and post-transcriptional modifications of Sf9 cells and tissue cells, which might alter the accessibility of the epitope to the primary α5-antibody. In this case, the tissue sample concentrations would also be lowered as a result of using relative standard curves produced from Sf9 cell derived NKA. The stark difference in detectability of rat NKA from recombinantly expressed Sf9 cells and from kidney tissue, both of which consist of the rat isoform α1 (ATP1A1), supports this hypothesis.

The NKA activity assays, which were run based on protein concentrations determined by the different quantification methods, helped to serve as an additional robustness assessment of the methods. The activity data of Sf9 cell-derived NKA showed consistently low variation across all animal species when the NKA activity assays were based on ELISA-determined protein concentrations (Fig. 8). The NKA activity assays based on concentrations determined by the three conventional methods exhibited different levels of variation across animal species—meaning that some animal species exhibited low variation in NKA activities while others exhibited high variation (Fig. 8). Due to these overlaps, it was not possible to detect a statistical difference between NKA activity variation in ELISA-based and other quantification method-based data. According to the ELISA, the determined NKA concentrations of protein samples derived from Sf9 cells were quite similar across the three biological replicates of each animal species. The conventional methods determined protein concentrations that were considerably lower, but, as with ELISA, they differ only slightly across biological replicates. This, in turn, resulted in lower variation in NKA activities for the conventional methods. The even NKA expression across biological replicates, as seen on the western blots (Fig. S2), further corroborate why no statistical difference was detected in the NKA activity variation between the four protein quantification methods. If NKA expression levels were uneven across biological replicates, then we would have expected to see more variation in the resulting activity data based on the conventional methods.

Of note, we found that the NKA activity assays of tissue-derived samples exhibited very high variation across the board. This was likely due to the absorbance values measured for the activity assays exceeding the linear range of the P_i_ standard curve, making accurate quantification of phosphate released from the hydrolysis of ATP impossible. There are two possible explanations for this outcome. The first is that NKAs derived from tissue have a higher catalytic activity than those expressed recombinantly. When the NKA activity is very high, then considerably less protein would be needed in the NKA activity assay to fall within the detectable range of the standard curve. Another explanation is that all four quantification methods, including the ELISA, severely underestimated the total NKA concentrations in tissue-derived samples, as was discussed previously. When the NKA concentration is underestimated, then more protein is used in the NKA activity assay, driving the NKA activity levels beyond the linear range of the standard curve. Despite numerous rounds of dilutions, we were never able to capture activity levels that fell within the standards range for our tissue-derived samples.

As part of the functional validation of the ELISA, several negative controls were run to test non-specific background noise in the ELISA. The negative control containing only NKA and the secondary HRP-antibody (“no α5-antibody”) showed a slightly higher absorbance than the detection threshold (Fig. 5). This minimal background noise most likely results from HRP-antibodies that remain in the wells after washing. For this reason, the number of wash steps following HRP-antibody incubation is higher than the rest of the washing steps. The absorbances of the other negative controls containing no HRP-conjugated secondary antibody or no protein sample did not exceed the detection threshold, indicating that TMB and the stopping solution do not stain non-specifically and that non-specific binding of the antibodies to the wells is not an issue in this ELISA.

Another important validation step involved confirming by western blot and ELISA that lyophilization does not alter the detectability of NKA by the antibodies used in the ELISA, thereby ensuring that the standards will not be detected differently than the fresh protein samples. Verifying that lyophilized cell membrane isolates can be used to produce relative protein standards is important considering how different the relative NKA standard curves can look for different animal species (Fig. 4). Without producing a standard curve for each animal species, the determined NKA concentrations would be biased towards the detectability of the protein’s animal species origin and thus significantly over-or underestimate NKA content in different samples. These interspecies differences are due to the α5-antibody binding at different efficiencies with different animal species (Fig. 4; Fig. S2).

Overall, our results indicate that the ELISA is the most suitable quantification method for the NKA, and most likely other large transmembrane proteins as well. The ELISA validated here can be easily adapted to other transmembrane proteins by exchanging the primary and, if necessary, the secondary

antibody. Moreover, it is straightforward to prepare relative protein standards that can be adapted to a wide taxonomic range and multiple sample types. This method eliminates the need for commercially available ELISA kits, which are more limited in these regards and can be quite costly.

## Acknowledgements

We thank M. Herbertz and V. Wagschal for their assistance in the laboratory. Funding: This study was supported by grants to SD from the Deutsche Forschungsgemeinschaft (Do 517/10-1) and to SM from the Alexander von Humboldt Foundation (Mohammadi 2018) and the National Institutes of Health (F32–HL149172).

