## Supplementary Information for "Evaluating the efficacy of protein quantification methods on membrane proteins"

**Table S1.** All materials and resources used in this study are included with sources and, when available, identifier number.

| **Reagent or Resource** | **Source** | **Identifier** |
|---|---|---|
| **Cell culture** | | |
| Insect-Xpress medium | Lonza, Walkersville, MD, USA | Cat#BE12-730P10 |
| Gentamycin | Roth, Karlsruhe, Germany | Cat#0233.1 |
| PEI MAX | Polysciences Inc., Warrington, PA, USA | CAS#49553-93-7 |
| **Biological samples** | | |
| Black rats | www.frostmaus.de | Cat#755190177-5 |
| Large milkweed bugs | AG Dobler, University of Hamburg, Germany | |
| Na, K-ATPase from porcine (Adenosine 5′-Triphosphatase from porcine cerebral cortex) (lyophilized) | Merck KGaA, Darmstadt, Germany | CAS#9000-83-3 |
| **Technical devices** | | |
| Ultrasonic homogenizer Sonopuls 2070 | Bandelin Electronic Company, Berlin, Germany | |
| Centrifuge 5840 R | Eppendorf AG, Hamburg, Germany | |
| Ultra-Centrifuge L-80 | Beckmann Coulter GmbH, Krefeld, Germany | |
| Freeze dryer Alpha 1-2 LD plus | Martin Christ Gefriertrocknungsanlagen GmbH, Osterode am Harz, Germany | |
| Ultrasonic homogenizer Omni Sonic Ruptor 400 | Omni International Inc., Kennesaw, GA, USA | |
| Cuvette spectrophotometer Ultrospec 2100 pro | Amersham Biosciences Europe GmbH, Freiburg im Breisgau, Germany | |
| Microplate shaker BioShake iQ | Quantifoil Instruments, Jena, Germany | |
| Microplate absorbance reader Bio-Rad Model 680 Bio-Rad iMark$^{TM}$ (spectrophotometers and software package) | **Bio-Rad Laboratories GmbH,** Feldkirchen, Germany | |
| Scanner Canon 9000F Mark II | Canon Europa N.V., Amsterdam, Netherlands | |
| **Commercial assay kits and material** | | |
| HPLC grade water ROTIPURAN® p.a., ACS water | Roth, Karlsruhe, Germany | Cat#HN68.2 |

| | | |
|---|---|---|
| Bovine serum albumin (BSA) | Roth, Karlsruhe, Germany | CAS#90604-29-8 |
| Coomassie Bradford reagent | Merck KGaA, Darmstadt, Germany | Cat#B6916 |
| Polystyrene cuvettes | SARSTEDT AG & Co. KG, Nümbrecht, Germany | Cat#67.742 |
| Modified Lowry protein assay kit | Thermo Fisher Scientific Inc., Rockford, IL, USA | Cat#23240 |
| 96-well polystyrene flat-well plate | SARSTEDT AG & Co. KG, Nümbrecht, Germany | Cat#82.1581 |
| Bicinchoninic acid protein assay kit | Thermo Fisher Scientific Inc., Rockford, IL, USA | Cat#23227 |
| Nitrocellulose membrane | Roth, Karlsruhe, Germany | Cat#HP42.1 |
| BlueBlock PF | SERVA Electrophoresis GmbH, Heidelberg, Germany | Cat#42591.01 |
| 4-Chloro-1-naphthol | Merck KGaA, Darmstadt, Germany | Cat#C8890 |
| Tween® 20 | Merck KGaA, Darmstadt, Germany | CAS#9005-64-5 |
| 3,3’,5,5’-tetramethylbenzidine (TMB) | Thermo Fisher Scientific Inc., Rockford, IL, USA | Cat#N301 |
| Parafilm® | Merck KGaA, Darmstadt, Germany | Cat#P7793 |
| Ouabain octahydrate, 96% | Thermo Fisher Scientific Inc., Rockford, IL, USA | CAS#11018-89-6 |
| Adenosin-5-triphosphat Bis-(Tris)-salt hydrate (ATP) | Merck KGaA, Darmstadt, Germany | CAS#102047-34-7 |
| **Antibodies** | | |
| α5-antibody (primary antibody) | Developmental Studies Hybridoma Bank, University of Iowa, Iowa City, IA, USA | RRID: AB_2166869 |
| HRP-antibody (secondary antibody conjugated with horseradish peroxidase) | Dianova, Hamburg, Germany | RRID: AB_2617176 |
| **Software** | | |
| ImageJ | https://imagej.net/software/imagej/ | ImageJ 1.53k |
| R | https://www.r-project.org/ | R version 4.1.1 |

**Table S2.** Addgene plasmid numbers, GenBank accession numbers, and CDS lengths for the ATP1A1 and ATP1B1 sequences used in the vectors of the six selected animal species. One plasmid (*Oncopeltus fasciatus*) was produced by S Dalla in [1] and is available upon request.

| Animal species | Addgene plasmid number | α-subunit gene | Β-subunit gene |
|---|---|---|---|
| *Rattus norvegicus* | 187416 | X05882<br>3072 bp | NM013113.2<br>915 bp |
| *Rhabdophis subminiatus* | 187418 | MT928191<br>3069 bp | ON168934<br>915 bp |
| *Tupinambis teguixin* | 187421 | MT928189<br>3069 bp | ON168937<br>918 bp |
| *Struthio camelus* | 187422 | XM009675281<br>3060 bp | XM009675170<br>747 bp |
| *Leptodactylus macrosternum* | 167172 | MT396189<br>3045 bp | SRR11583987<br>915 bp |
| *Oncopeltus fasciatus* | - | OW028346<br>3024 bp | OW028347<br>993 bp |

1. Herbertz M, Dalla S, Wagschal V, Turjalei R, Heiser M, Dobler S. 2023 Co-evolutionary escalation led to differentially adapted paralogs of an insect's Na,K-ATPase optimizing resistance to host plant toxins. *Molecular Ecology*

**Table S3.** Western blot and ELISA raw data for non-lyophilized and lyophilized protein samples. Western blot samples were run on two gels, with brown rat, common ostrich, large milkweed bug, and red-necked keelback snake on one gel, and gold tegu, Miranda's white-lipped frog, and uninfected Sf0 cells on a second. Densities of Na,K-ATPase α-subunit bands were determined by the areas under the curves in ImageJ (see plots created with ImageJ in Fig. S2 and membrane scans in Fig. 3). For the ELISA, the average of two technical replicates (M1 and M2) run on one plate was used for subsequent analyses. All protein samples were diluted 1:2000 prior to running them on the ELISA.

| **Species** | **Treatment** | **Area under curve** |
|---|---|---|
| untransfected Sf9 cells | non-lyophilized | 1052 |
| | lyophilized | 1539 |
| brown rat | non-lyophilized | 4488 |
| | lyophilized | 3972 |
| common ostrich | non-lyophilized | 3487 |
| | lyophilized | 4582 |
| large milkweed bug | non-lyophilized | 1968 |
| | lyophilized | 1560 |
| gold tegu | non-lyophilized | 3985 |
| | lyophilized | 2942 |
| Miranda's white-lipped frog | non-lyophilized | 3419 |
| | lyophilized | 3937 |
| red-necked keelback snake | non-lyophilized | 3688 |
| | lyophilized | 3711 |

| **ELISA** | | **Absorbance [490 nm]** | | |
|---|---|---|---|---|
| **Species** | **Treatment** | **M1** | **M2** | **average** |
| untransfected Sf9 cells | non-lyophilized | 0.229 | 0.248 | 0.239 |
| | lyophilized | 0.201 | 0.256 | 0.229 |
| brown rat | non-lyophilized | 0.576 | 0.570 | 0.573 |
| | lyophilized | 0.598 | 0.602 | 0.600 |
| common ostrich | non-lyophilized | 0.911 | 0.969 | 0.940 |
| | lyophilized | 1.004 | 0.993 | 0.999 |
| large milkweed bug | non-lyophilized | 0.641 | 0.693 | 0.667 |
| | lyophilized | 0.893 | 0.909 | 0.901 |
| gold tegu | non-lyophilized | 0.723 | 0.705 | 0.714 |
| | lyophilized | 0.788 | 0.789 | 0.789 |
| Miranda's white-lipped frog | non-lyophilized | 0.694 | 0.691 | 0.693 |
| | lyophilized | 0.816 | 0.788 | 0.802 |
| red-necked keelback snake | non-lyophilized | 0.479 | 0.498 | 0.489 |
| | lyophilized | 0.509 | 0.461 | 0.485 |

**Table S4.** Modified Lowry protein assay raw data for the BSA standard curve, protein samples, and the corresponding downstream Na, K-ATPase activity based on Lowry assay concentrations. The average of two technical replicates (M1 and M2) were used to calculate the final concentration for each sample.

| **BSA standard** | | | |
|---|---|---|---|
| **Concentration [µg/mL]** | **Absorbance [655 nm]** | | |
| | **M1** | **M2** | **average** |
| 0 | 0.123 | 0.078 | 0.101 |
| 1 | 0.093 | 0.060 | 0.077 |
| 5 | 0.067 | 0.061 | 0.064 |
| 25 | 0.184 | 0.172 | 0.178 |
| 125 | 0.212 | 0.195 | 0.204 |
| 250 | 0.421 | 0.462 | 0.442 |
| 500 | 0.701 | 0.708 | 0.705 |
| 750 | 0.848 | 0.917 | 0.883 |
| 1000 | 1.155 | 1.201 | 1.178 |
| 1500 | 1.381 | 1.371 | 1.376 |
| 2000 | 1.636 | 1.636 | 1.636 |

| **Lowry assay of Sf9 cells-derived samples** | | | | | | |
|---|---|---|---|---|---|---|
| **Species** | **Biological replicates** | **Absorbance [655 nm]** | | | **Calculated concentration [µg/mL]** | **Na, K-ATPase activity [nmol Pi/(mg protein*min)]** |
| | | **M1** | **M2** | **average** | | |
| **uninfected Sf9 cells** | A | 0.963 | 0.957 | 0.960 | 9550 | - |
| **brown rat** | A | 1.161 | 1.140 | 1.151 | 11667 | 7.78 |
| | B | 1.146 | 1.147 | 1.147 | 11622 | 10.24 |
| | C | 1.264 | 1.334 | 1.299 | 13317 | 8.96 |
| **common ostrich** | A | 1.084 | 0.708 | 0.896 | 8839 | 33.20 |
| | B | 1.486 | 1.431 | 1.459 | 15089 | 17.78 |
| | C | 1.929 | 1.515 | 1.722 | 18017 | 19.26 |
| **large milkweed bug** | A | 1.585 | 1.652 | 1.619 | 16867 | 38.32 |
| | B | 1.402 | 1.311 | 1.357 | 13956 | 35.10 |
| | C | 1.214 | 1.276 | 1.245 | 12717 | 38.84 |
| **gold tegu** | A | 1.024 | 1.071 | 1.048 | 10522 | 10.51 |
| | B | 1.018 | 0.981 | 1.000 | 9989 | 9.99 |
| | C | 0.853 | 0.600 | 0.727 | 6956 | 10.41 |
| **Miranda's white-lipped frog** | A | 1.428 | 1.267 | 1.348 | 13856 | - |
| | B | 1.126 | 1.113 | 1.120 | 11322 | - |
| | C | 0.256 | 0.241 | 0.249 | 1644 | - |
| **red-necked keelback snake** | A | 1.383 | 1.347 | 1.365 | 14050 | 8.09 |
| | B | 1.570 | 1.523 | 1.547 | 16067 | 5.98 |
| | C | 1.491 | 1.392 | 1.442 | 14900 | 2.42 |
| **Lowry assay of tissue-derived samples** | | | | | | |
| **black rat brain** | A | 0.996 | 0.976 | 0.986 | 9839 | 66.16 |
| | B | 0.884 | 0.869 | 0.877 | 8622 | 21.58 |
| | C | 1.151 | 1.127 | 1.139 | 11539 | 78.57 |
| **black rat kidney** | A | 0.452 | 0.436 | 0.444 | 4033 | 92.09 |
| | B | 0.347 | 0.326 | 0.337 | 2839 | 92.75 |
| | C | 0.300 | 0.285 | 0.293 | 2350 | 85.58 |
| **large milkweed bug brain** | A | 0.638 | 0.626 | 0.632 | 591 | - |
| **porcine brain** | A | 0.800 | 0.791 | 0.796 | 772 | - |

**Table S5.** Bicinchoninic acid (BCA) protein assay raw data for the BSA standard curve, protein samples, and the corresponding downstream Na, K-ATPase activity based on BCA assay concentrations. The average of two technical replicates (M1 and M2) were used to calculate the final concentration for each sample.

| **BSA standard** | | | |
|---|---|---|---|
| **Concentration [µg/mL]** | **Absorbance [550 nm]** | | |
| | **M1** | **M2** | **average** |
| 0 | 0.085 | 0.080 | 0.083 |
| 1 | 0.086 | 0.085 | 0.086 |
| 5 | 0.091 | 0.092 | 0.092 |
| 25 | 0.133 | 0.117 | 0.125 |
| 125 | 0.348 | 0.341 | 0.345 |
| 250 | 0.520 | 0.553 | 0.537 |
| 500 | 0.916 | 0.935 | 0.926 |
| 750 | 1.111 | 1.197 | 1.154 |
| 1000 | 1.616 | 1.578 | 1.597 |
| 1500 | 2.020 | 2.083 | 2.052 |
| 2000 | 2.440 | 2.491 | 2.466 |

| **BCA assay of Sf9 cells-derived samples** | | | | | | |
|---|---|---|---|---|---|---|
| **Species** | **Biological replicates** | **Absorbance [550 nm]** | | | **Calculated concentration [µg/mL]** | **Na, K-ATPase activity [nmol Pi/(mg protein*min)]** |
| | | **M1** | **M2** | **average** | | |
| **uninfected Sf9 cells** | A | 1.139 | 1.103 | 1.121 | 7988 | - |
| **brown rat** | A | 1.300 | 1.233 | 1.267 | 9108 | 21.86 |
| | B | 1.302 | 1.320 | 1.311 | 9450 | 24.07 |
| | C | 1.453 | 1.704 | 1.579 | 11508 | 7.57 |
| **common ostrich** | A | 1.301 | 1.271 | 1.286 | 9258 | 25.52 |
| | B | 2.072 | 2.068 | 2.070 | 15288 | 17.84 |
| | C | 2.371 | 2.545 | 2.458 | 18273 | 26.18 |
| **large milkweed bug** | A | 2.071 | 2.013 | 2.042 | 15073 | 42.12 |
| | B | 1.505 | 1.541 | 1.523 | 11081 | 34.38 |
| | C | 1.502 | 1.508 | 1.505 | 10942 | 50.39 |
| **gold tegu** | A | 1.162 | 1.090 | 1.126 | 8027 | 10.89 |
| | B | 1.117 | 1.068 | 1.093 | 7769 | 7.88 |
| | C | 0.967 | 0.931 | 0.949 | 6665 | 5.60 |
| **Miranda's white-lipped frog** | A | 1.606 | 1.548 | 1.577 | 11496 | - |
| | B | 1.204 | 1.229 | 1.217 | 8723 | - |
| | C | 0.256 | 0.234 | 0.245 | 1250 | - |
| **red-necked keelback snake** | A | 1.516 | 1.508 | 1.512 | 10996 | 4.08 |
| | B | 1.799 | 1.753 | 1.776 | 13027 | 7.47 |
| | C | 1.567 | 1.644 | 1.606 | 11715 | 10.03 |
| **BCA assay of tissue-derived samples** | | | | | | |
| **black rat brain** | A | 1.292 | 1.305 | 1.299 | 9354 | 65.98 |
| | B | 1.688 | 1.438 | 1.563 | 8012 | 15.84 |
| | C | 1.356 | 1.504 | 1.430 | 10365 | 77.74 |
| **black rat kidney** | A | 0.418 | 0.406 | 0.412 | 2531 | 98.25 |
| | B | 0.324 | 0.346 | 0.335 | 1938 | 99.04 |
| | C | 0.271 | 0.281 | 0.276 | 1485 | 86.99 |
| **large milkweed bug brain** | A | 0.776 | 0.808 | 0.792 | 546 | - |
| **porcine brain** | A | 0.857 | 0.824 | 0.841 | 583 | - |

**Table S6.** Coomassie Bradford protein assay raw data for the BSA standard curve, protein samples, and the corresponding downstream Na, K-ATPase activity based on Bradford assay concentrations. The average of two technical replicates (M1 and M2) were used to calculate the final concentration for each sample.

| **BSA standard** | | | |
|---|---|---|---|
| **Concentration [µg/mL]** | **Absorbance [595 nm]** | | |
| | **M1** | **M2** | **average** |
| 0 | 0.000 | 0.000 | 0.000 |
| 2 | 0.097 | 0.067 | 0.082 |
| 4 | 0.191 | 0.151 | 0.171 |
| 6 | 0.241 | 0.22 | 0.231 |
| 8 | 0.297 | 0.276 | 0.287 |
| 10 | 0.353 | 0.348 | 0.351 |

| **Bradford assay of Sf9 cells-derived samples** | | | | | | |
|---|---|---|---|---|---|---|
| **Species** | **Biological replicates** | **Absorbance [595 nm]** | | | **Calculated concentration [µg/mL]** | **Na, K-ATPase activity [nmol Pi/(mg protein*min)]** |
| | | **M1** | **M2** | **average** | | |
| **uninfected Sf9 cells** | A | 0.199 | 0.088 | 0.144 | 7863 | - |
| **brown rat** | A | 0.190 | 0.233 | 0.212 | 11589 | 10.37 |
| | B | 0.182 | 0.225 | 0.204 | 11151 | 8.65 |
| | C | 0.250 | 0.236 | 0.243 | 13315 | 7.23 |
| **common ostrich** | A | 0.208 | 0.230 | 0.219 | 12000 | 27.77 |
| | B | 0.284 | 0.151 | 0.218 | 11918 | 18.67 |
| | C | 0.214 | 0.319 | 0.267 | 14603 | 3.77 |
| **large milkweed bug** | A | 0.286 | 0.301 | 0.294 | 16082 | 36.17 |
| | B | 0.257 | 0.080 | 0.169 | 9233 | 34.48 |
| | C | 0.236 | 0.198 | 0.217 | 11890 | 43.57 |
| **gold tegu** | A | 0.177 | 0.199 | 0.188 | 10301 | 8.40 |
| | B | 0.193 | 0.192 | 0.193 | 10548 | 6.29 |
| | C | 0.232 | 0.190 | 0.211 | 11562 | 7.78 |
| **Miranda's white-lipped frog** | A | 0.170 | 0.245 | 0.208 | 11370 | - |
| | B | 0.229 | 0.238 | 0.234 | 12795 | - |
| | C | 0.101 | -0.051 | 0.025 | 1370 | - |
| **red-necked keelback snake** | A | 0.278 | 0.262 | 0.270 | 14795 | 6.02 |
| | B | 0.402 | 0.284 | 0.343 | 18795 | 4.12 |
| | C | 0.207 | 0.276 | 0.242 | 13233 | 6.36 |
| **Bradford assay of tissue cells-derived samples** | | | | | | |
| **black rat brain** | A | 0.072 | 0.062 | 0.067 | 3671 | 29.95 |
| | B | 0.067 | 0.058 | 0.063 | 3425 | 50.53 |
| | C | 0.104 | 0.081 | 0.093 | 5068 | 56.79 |
| **black rat kidney** | A | 0.167 | 0.164 | 0.166 | 9068 | 100.40 |
| | B | 0.099 | 0.097 | 0.098 | 5370 | 93.37 |
| | C | 0.115 | 0.115 | 0.115 | 6301 | 78.01 |
| **large milkweed bug brain** | A | 0.150 | 0.333 | 0.242 | 1321 | - |
| **porcine brain** | A | 0.195 | 0.314 | 0.255 | 1392 | - |

**Table S7.** ELISA buffer recipes.

| **Name** | **Ingredients** |
|---|---|
| Phosphate-buffered saline (PBS) | 8.0 g NaCl<br>0.2 g $KH_2PO_4$<br>1.1 g $Na_2HPO_4$<br>0.2 g KCl<br><br>ad 1 L Millipore water |
| coating buffer | 0.841 g $NaHCO_3$<br><br>ad 100 mL PBS<br>adjust pH to 9.5 with KOH |
| washing buffer | 0.5 mL Tween®20<br><br>in 1 L PBS |
| blocking buffer | 1 g BSA<br>200 µL Tween®20<br><br>in 100 mL PBS |
| stopping solution | 5.556 mL 18 M $H_2SO_4$<br><br>ad 194.444 mL Millipore water |

**Table S8.** Absorbance values for ELISA's negative controls. A relative randomly selected sample from our transfected Sf9 cell derived samples (gold tegu replicate A; Table S9) was selected for the negative controls that contained protein (i.e., no α5-antibody and no HRP-antibody). The average of two technical replicates (M1 and M2) run on one plate was used for subsequent graphing (Fig. 5).

| | **Absorbance [490 nm]** | | |
|---|---|---|---|
| **Negative control** | **M1** | **M2** | **average** |
| Transfected Sf9 cells | 0.826 | 0.840 | 0.833 |
| untransfected Sf9 cells | 0.333 | 0.376 | 0.355 |
| no α5-antibody | 0.070 | 0.077 | 0.074 |
| no HRP-antibody | 0.039 | 0.042 | 0.041 |
| no protein | 0.036 | 0.036 | 0.036 |
| no protein, no antibodies | 0.033 | 0.035 | 0.034 |

**Table S9.** ELISA raw data for relative Na, K-ATPase standard curves (grey), protein samples (light grey), and the corresponding $Na^{+}$, $K^{+}$-ATPase activity based on ELISA concentrations (white). The average of two technical replicates (M1 and M2) were used to calculate the final concentration for each sample.

| **Species** | **Concentration [µg/mL]** | **Absorbance [490 nm]** | | | **Biological replicates** | **Absorbance [490 nm]** | | | **Calculated concentration [µg/mL]** | **Na, K-ATPase activity [nmol Pi/(mg protein*min)]** |
|---|---|---|---|---|---|---|---|---|---|---|
| | | **M1** | **M2** | **average** | | **M1** | **M2** | **average** | | |
| **brown rat** | 0 | 0.036 | 0.041 | 0.039 | **A** (1:2000) | 0.629 | 0.632 | 0.631 | 16109 | 3.42 |
| | 0.625 | 0.094 | 0.098 | 0.096 | **B** (1:2000) | 0.583 | 0.619 | 0.601 | 15306 | 7.85 |
| | 0.9375 | 0.134 | 0.144 | 0.139 | **C** (1:2000) | 0.637 | 0.662 | 0.650 | 16626 | 6.19 |
| | 1.25 | 0.143 | 0.146 | 0.145 | **Rat brain A** (1:2000) | 0.611 | 0.626 | 0.619 | 15782 | - |
| | 1.875 | 0.202 | 0.198 | 0.200 | **Rat brain B** (1:2000) | 0.515 | 0.518 | 0.517 | 13007 | - |
| | 2.5 | 0.287 | 0.272 | 0.280 | **Rat brain C** (1:2000) | 0.481 | 0.510 | 0.496 | 12435 | - |
| | 3.75 | 0.373 | 0.352 | 0.363 | **Rat kidney A** (1:2000) | 0.244 | 0.246 | 0.245 | 5755 | - |
| | 5 | 0.435 | 0.421 | 0.428 | **Rat kidney B** (1:2000) | 0.200 | 0.204 | 0.202 | 4585 | - |
| | 7.5 | 0.603 | 0.596 | 0.600 | **Rat kidney C** (1:2000) | 0.163 | 0.180 | 0.172 | 3755 | - |
| | 10 | 0.703 | 0.722 | 0.713 | | | | | | |
| | 15 | 0.789 | 0.808 | 0.799 | | | | | | |
| | 20 | 0.86 | 0.888 | 0.874 | | | | | | |
| | 30 | 0.912 | 0.945 | 0.929 | | | | | | |
| | 60 | 0.914 | 0.932 | 0.923 | | | | | | |

| | | | | | | | | | | |
|---|---|---|---|---|---|---|---|---|---|---|
| **common ostrich** | 0 | 0.036 | 0.036 | 0.036 | **A** (1:6000) | 0.829 | 0.910 | 0.870 | 135569 | 4.98 |
| | 0.625 | 0.084 | 0.069 | 0.077 | **B** (1:6000) | 0.850 | 0.823 | 0.837 | 130203 | 4.98 |
| | 0.9375 | 0.102 | 0.096 | 0.099 | **C** (1:6000) | 0.941 | 0.925 | 0.933 | 145894 | 8.06 |
| | 1.25 | 0.126 | 0.107 | 0.117 | | | | | | |
| | 1.875 | 0.149 | 0.148 | 0.149 | | | | | | |
| | 2.5 | 0.204 | 0.196 | 0.200 | | | | | | |
| | 3.75 | 0.274 | 0.274 | 0.274 | | | | | | |
| | 5 | 0.371 | 0.374 | 0.373 | | | | | | |
| | 7.5 | 0.38 | 0.389 | 0.385 | | | | | | |

| | | | | |
|---|---|---|---|---|
| | 10 | 0.545 | 0.531 | 0.538 |
| | 15 | 0.749 | 0.795 | 0.772 |
| | 20 | 0.832 | 0.833 | 0.833 |
| | 30 | 0.893 | 0.907 | 0.900 |
| | 60 | 1.017 | 1.065 | 1.041 |

| | | | | | | | | | | |
|---|---|---|---|---|---|---|---|---|---|---|
| **large milkweed bug** | 0 | 0.034 | 0.036 | 0.035 | **A** (1:4000) | 0.889 | 0.868 | 0.879 | 62311 | 16.18 |
| | 0.625 | 0.096 | 0.101 | 0.099 | **B** (1:4000) | 0.809 | 0.802 | 0.806 | 56913 | 12.90 |
| | 0.9375 | 0.130 | 0.121 | 0.126 | **C** (1:4000) | 0.858 | 0.861 | 0.860 | 60906 | 13.42 |
| | 1.25 | 0.153 | 0.155 | 0.154 | **Milkweed bug brain A** (1:2000) | 0.073 | 0.072 | 0.073 | 1386 | - |
| | 1.875 | 0.196 | 0.192 | 0.194 | | | | | | |
| | 2.5 | 0.248 | 0.255 | 0.252 | | | | | | |
| | 3.75 | 0.316 | 0.316 | 0.316 | | | | | | |
| | 5 | 0.453 | 0.455 | 0.454 | | | | | | |
| | 7.5 | 0.557 | 0.574 | 0.566 | | | | | | |
| | 10 | 0.724 | 0.728 | 0.726 | | | | | | |
| | 15 | 0.801 | 0.833 | 0.817 | | | | | | |
| | 20 | 0.906 | 0.978 | 0.942 | | | | | | |
| | 30 | 0.969 | 1.027 | 0.998 | | | | | | |
| | 60 | 1.082 | 1.117 | 1.100 | | | | | | |

| | | | | | | | | | | |
|---|---|---|---|---|---|---|---|---|---|---|
| **gold tegu** | 0 | 0.037 | 0.035 | 0.036 | **A** (1:2000) | 0.826 | 0.840 | 0.833 | 14186 | 6.88 |
| | 0.625 | 0.177 | 0.168 | 0.173 | **B** (1:2000) | 0.858 | 0.885 | 0.872 | 14871 | 5.46 |
| | 0.9375 | 0.219 | 0.220 | 0.220 | **C** (1:2000) | 0.894 | 0.938 | 0.916 | 15663 | 6.09 |
| | 1.25 | 0.272 | 0.263 | 0.268 | | | | | | |
| | 1.875 | 0.369 | 0.368 | 0.369 | | | | | | |
| | 2.5 | 0.521 | 0.529 | 0.525 | | | | | | |
| | 3.75 | 0.609 | 0.601 | 0.605 | | | | | | |
| | 5 | 0.763 | 0.765 | 0.764 | | | | | | |
| | 7.5 | 0.86 | 0.847 | 0.854 | | | | | | |
| | 10 | 0.939 | 0.944 | 0.942 | | | | | | |
| | 15 | 0.973 | 0.991 | 0.982 | | | | | | |
| | 20 | 1.083 | 1.094 | 1.089 | | | | | | |
| | 30 | 1.056 | 1.086 | 1.071 | | | | | | |
| | 60 | 1.089 | 1.118 | 1.104 | | | | | | |

| | | | | | | | | | | |
|---|---|---|---|---|---|---|---|---|---|---|
| | 0 | 0.034 | 0.036 | 0.035 | **A** (1:2000) | 0.870 | 0.879 | 0.875 | 16874 | - |

| | | | | | | | | | | |
|---|---|---|---|---|---|---|---|---|---|---|
| **Miranda's white-lipped frog** | 0.625 | 0.143 | 0.136 | 0.140 | **B** (1:2000) | 0.770 | 0.792 | 0.781 | 14995 | - |
| | 0.9375 | 0.189 | 0.198 | 0.194 | **C** (1:2000) | 0.053 | 0.050 | 0.052 | 332 | - |
| | 1.25 | 0.218 | 0.216 | 0.217 | | | | | | |
| | 1.875 | 0.271 | 0.280 | 0.276 | | | | | | |
| | 2.5 | 0.387 | 0.363 | 0.375 | | | | | | |
| | 3.75 | 0.504 | 0.500 | 0.502 | | | | | | |
| | 5 | 0.621 | 0.636 | 0.629 | | | | | | |
| | 7.5 | 0.786 | 0.786 | 0.786 | | | | | | |
| | 10 | 0.918 | 0.870 | 0.894 | | | | | | |
| | 15 | 1.007 | 1.011 | 1.009 | | | | | | |
| | 20 | 1.033 | 1.044 | 1.039 | | | | | | |
| | 30 | 1.041 | 1.067 | 1.054 | | | | | | |
| | 60 | 1.028 | 1.041 | 1.035 | | | | | | |

| | | | | | | | | | | |
|---|---|---|---|---|---|---|---|---|---|---|
| **red-necked keelback snake** | 0 | 0.036 | 0.041 | 0.039 | **A** (1:2000) | 0.316 | 0.352 | 0.334 | 48443 | 1.49 |
| | 0.625 | 0.046 | 0.043 | 0.045 | **B** (1:2000) | 0.326 | 0.372 | 0.349 | 50902 | 3.15 |
| | 0.9375 | 0.043 | 0.043 | 0.043 | **C** (1:2000) | 0.362 | 0.397 | 0.380 | 55902 | 2.80 |
| | 1.25 | 0.066 | 0.047 | 0.057 | | | | | | |
| | 1.875 | 0.051 | 0.051 | 0.051 | | | | | | |
| | 2.5 | 0.075 | 0.073 | 0.074 | | | | | | |
| | 3.75 | 0.071 | 0.080 | 0.076 | | | | | | |
| | 5 | 0.109 | 0.122 | 0.116 | | | | | | |
| | 7.5 | 0.121 | 0.125 | 0.123 | | | | | | |
| | 10 | 0.173 | 0.200 | 0.187 | | | | | | |
| | 15 | 0.182 | 0.175 | 0.179 | | | | | | |
| | 20 | 0.343 | 0.353 | 0.348 | | | | | | |
| | 30 | 0.476 | 0.399 | 0.438 | | | | | | |
| | 60 | 0.72 | 0.759 | 0.740 | | | | | | |

| | | | | | | | | | | |
|---|---|---|---|---|---|---|---|---|---|---|
| **porcine** | 0 | 0.033 | 0.034 | 0.034 | 10 µg/mL | 0.338 | 0.348 | 0.343 | 16913 | - |
| | 0.625 | 0.039 | 0.040 | 0.040 | | | | | | |
| | 0.9375 | 0.057 | 0.051 | 0.054 | | | | | | |
| | 1.25 | 0.047 | 0.045 | 0.046 | | | | | | |
| | 1.875 | 0.064 | 0.059 | 0.062 | | | | | | |
| | 2.5 | 0.064 | 0.065 | 0.065 | | | | | | |
| | 3.75 | 0.104 | 0.095 | 0.100 | | | | | | |
| | 5 | 0.106 | 0.102 | 0.104 | | | | | | |
| | 7.5 | 0.175 | 0.161 | 0.168 | | | | | | |
| | 10 | 0.217 | 0.202 | 0.210 | | | | | | |

| | | | | |
|---|---|---|---|---|
| | 15 | 0.290 | 0.327 | 0.309 |
| | 20 | 0.423 | 0.401 | 0.412 |
| | 30 | 0.510 | 0.553 | 0.532 |
| | 60 | 0.726 | 0.744 | 0.735 |

**Table S10.** P-values of post-hoc Dunn's test comparing the differences in the rank sums of the determined protein concentrations between the four protein quantification methods. Marked differences: ** $p < 0.05$ (significant)

| **Samples derived from Sf9 cells** | | **Adjusted p-value (Dunn's test)** |
|---|---|---|
| Lowry | BCA | 0.545 |
| Lowry | Bradford | 1 |
| Lowry | ELISA | 0.00119 ** |
| BCA | Bradford | 1 |
| BCA | ELISA | 0.000000374 ** |
| Bradford | ELISA | 0.00033 ** |
| **Samples derived from tissue isolates** | | |
| Lowry | BCA | 1 |
| Lowry | Bradford | 1 |
| Lowry | ELISA | 1 |
| BCA | Bradford | 1 |
| BCA | ELISA | 0.587 |
| Bradford | ELISA | 0.190 |

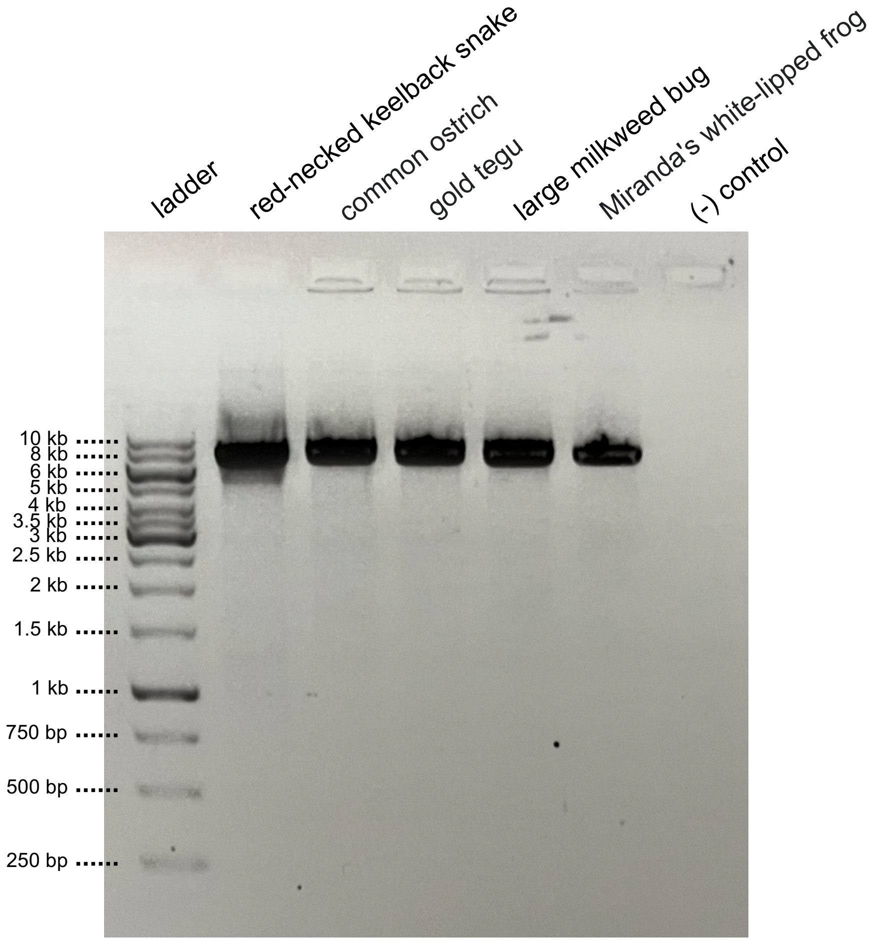

**Figure S1.** Agarose gel electrophoresis of recombinant bacmid DNA amplified with PCR to verify the transposition of the genes of interest into the bacmids. The expected total length of the PCR products with inserts is around 6600-6900 bp.

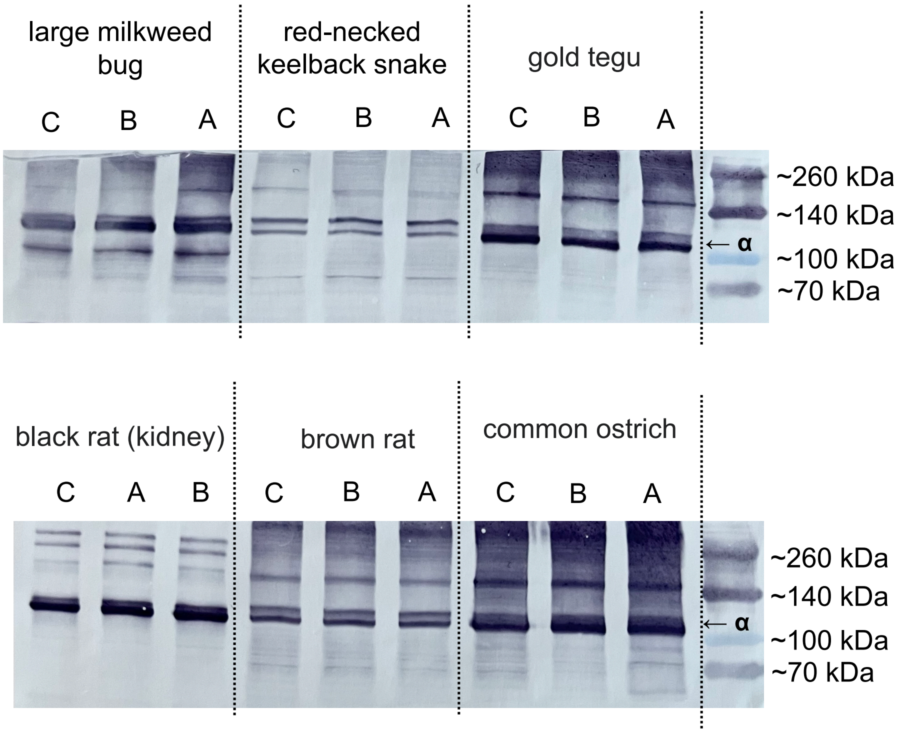

**Figure S2.** Western blot analysis of Na,K-ATPase (NKA) derived from Sf9 cell expression and from black rat kidney tissue. Black rat brain tissue samples were used up in the activity assays and therefore excluded from this western blot analysis. The 110 kDA α-subunits of the protein are stained with the α5 monoclonal antibody followed by a horseradish peroxidase conjugated goat anti-mouse antibody. Each panel shows the western blot of one gel, with the ladder on the right. Lanes show three biological replicates (A, B, and C) for each NKA type. 2.5 uL of protein sample was used in all western blots to visualize differences in NKA content across samples.

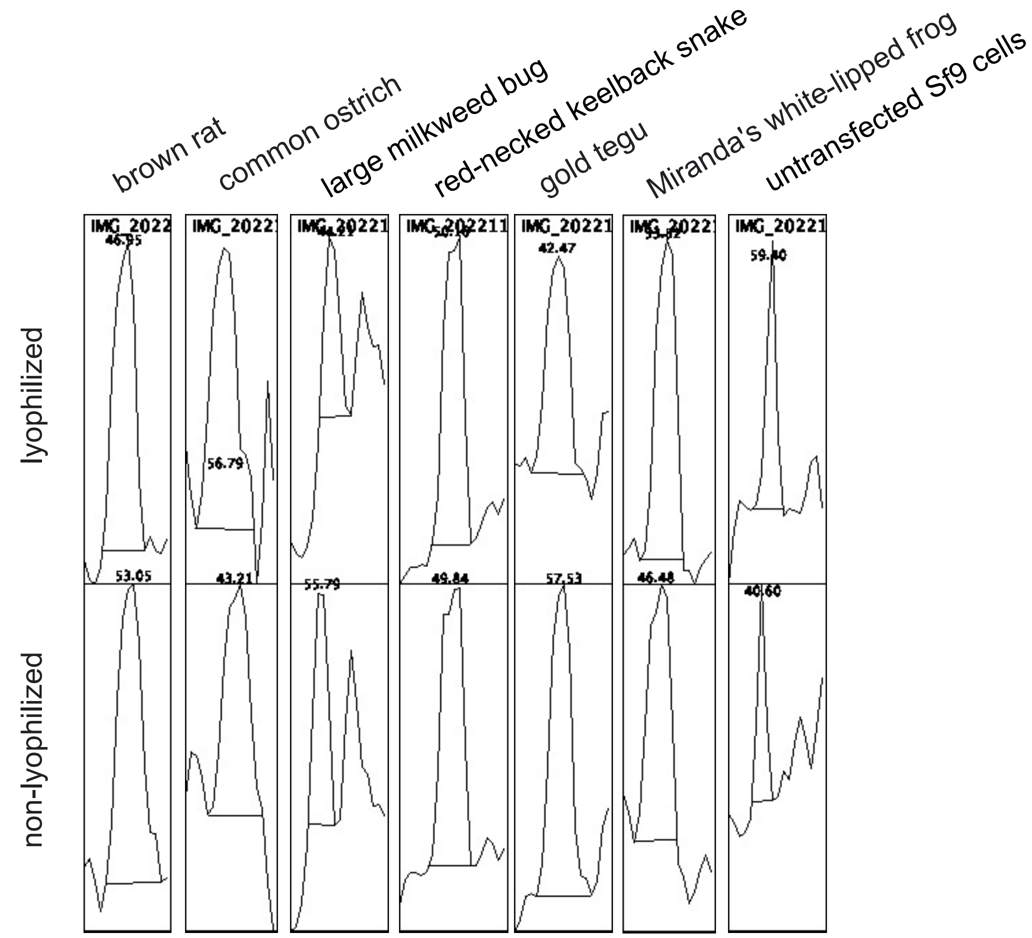

**Figure S3.** Western blot band density plots created with ImageJ to compare the areas under the curves of lyophilized and non-lyophilized protein samples.
